# A zebrafish model of Granulin deficiency reveals essential roles in myeloid cell differentiation

**DOI:** 10.1101/2020.07.23.217067

**Authors:** Clyde A. Campbell, Oksana Fursova, Xiaoyi Cheng, Elizabeth Snella, Abbigail McCune, Liangdao Li, Barbara Solchenberger, Bettina Schmid, Debashis Sahoo, Mark Morton, David Traver, Raquel Espín-Palazón

**Affiliations:** Department of Genetics, Development and Cell Biology. Iowa State University, 2213 Pammel Drive, Advance Teaching and Research Building 3003, Ames, IA, 50011, USA; Section of Cell and Developmental Biology. University of California at San Diego, 9500 Gilman Drive, Natural Sciences Building 6107, La Jolla, CA, 92093, USA; German Center for Neurodegenerative Diseases (DZNE), Munich, Germany; Department of Computer Science and Engineering. University of California at San Diego, La Jolla, CA, 92093, USA; College of Veterinary Medicine. Iowa State University, IA, USA

## Abstract

Granulin (*GRN*) is a pleiotropic protein involved in inflammation, wound healing, neurodegenerative disease, and tumorigenesis. These roles in human health have prompted research efforts to utilize Granulin in the treatment of rheumatoid arthritis, frontotemporal dementia, and to enhance wound healing. How granulin contributes to each of these diverse biological functions, however, remains largely unknown. Here, we have uncovered a new role for granulin during myeloid cell differentiation. Using a zebrafish model of granulin deficiency, we reveal that in the absence of granulin a (*grna*), myeloid progenitors are unable to terminally differentiate into neutrophils and macrophages during normal and emergency myelopoiesis. In addition, macrophages fail to recruit to the wound, resulting in abnormal healing. Our CUT&RUN experiments identify Pu.1, which together with Irf8 positively regulate *grna* expression. Importantly, we demonstrate functional conservation between the mammalian granulin and the zebrafish orthologue *grna*. Our findings uncover a previously unrecognized role for granulin during myeloid cell differentiation, opening a new field of study that has the potential to impact different aspects of the human health.

## Introduction

Neutrophils and macrophages differentiate from myeloid progenitors and are essential for clearing infections and promoting tissue repair. In addition, recent studies have elucidated a multitude of other functions beyond their classical inflammatory roles. For instance, we and others have demonstrated that neutrophils and macrophages are critical during hematopoietic stem cell (HSC) specification (Espin-Palazon *et al*, 2014; He *et al*, 2015; Li *et al*, 2014; Theodore *et al*, 2017; Travnickova *et al*, 2015). Numerous studies have shown that microglia, the tissue resident macrophages of the brain, are involved in neurodegenerative disease (Bachiller *et al*, 2018; Wynn *et al*, 2013). In addition, tumor-associated macrophages (TAMs) and neutrophils (TANs) can be key in contributing to tumor metastasis. The tremendous implications that neutrophils and macrophages have in tissue homeostasis and how their disruption leads to human disease have put these cells in the spotlight of many recent investigations. The identification of new molecular regulators of myeloid cell differentiation thus has the potential to broadly impact human health.

Granulin (*GRN*) is a protein of pleiotropic function that contains several cysteine-rich motifs unique to this molecule (Bhandari *et al*, 1992). It was first isolated from leukocytes (Bateman *et al*, 1990), and It is known to regulate inflammation, wound healing, tissue growth, and it is involved in neurodegenerative diseases, lipofuscinosis, and tumorigenesis (Bateman *et al*, 2018). Although its cognate receptor has not been identified, granulin can bind to a wide variety of receptors, including TNF receptors, Ephrin type-A receptor 2 (EphA2), Notch, Toll-like receptor 9 (TLR9), LDL Receptor Related Protein 1 (LRP1), and sortilin1 (Chitramuthu *et al*, 2017). Consequently, Granulin can modulate diverse signaling pathways such as NFkB, WNT, MAPK/ERK, PI3K/Akt, and FAK (Alquezar *et al*, 2016; de la Encarnacion *et al*, 2016; He *et al*, 2002; Hwang *et al*, 2013; Lu & Serrero, 2001; Tian *et al*, 2016; Zanocco-Marani *et al*, 1999).

Many studies support a critical role for granulin during infection and inflammation. While granulin can function as an anti-inflammatory factor, it can also exert a pro-inflammatory role. *GRN* knockout mice fail to clear *Listeria monocytogenes*, and their macrophages express high levels of pro-inflammatory cytokines. However, it is surprising that only a few macrophages were found in infected organs (Yin *et al*, 2010). Moreover, Granulin functions as an antagonist of TNF signaling, playing a critical role in the pathogenesis of inflammatory diseases (Zhu *et al*, 2002), and has been reported as a promising therapeutic target for inflammatory diseases including rheumatoid arthritis, psoriasis, and osteoarthritis (Farag *et al*, 2019; Liu, 2011; Tang *et al*, 2011; Wei *et al*, 2017).

Several studies have demonstrated that Granulin facilitates wound healing (for review, see Jian and colleagues (Jian *et al*, 2013)). Administration of GRN to fresh wounds increased the cell counts of neutrophils, macrophages, and fibroblasts, which collectively facilitated healing (He *et al*, 2003). In addition, it has been shown that macrophages produce granulin as a key regulatory factor in the processes of inflammation and wound healing (Yin *et al*., 2010).

Mammals encode a single granulin gene (*GRN*), expressed in most tissues (Bhandari *et al*, 1993; Daniel *et al*, 2000). Although recent work indicates that granulin regulates the formation of regulatory T lymphocytes (Kwack & Lee, 2017; Wei *et al*, 2014), human hematopoietic cells from the myeloid lineage contain the highest levels of granulin transcripts, being one of the most abundant transcripts in macrophages (Chantry *et al*, 1998) and monocyte-derived dendritic cells (Hashimoto *et al*, 1999). Although previous work showed a positive correlation between myeloid cell maturation and *GRN* expression (Ong *et al*, 2006), a potential role for *GRN* in myelopoiesis *in vivo* has not been reported.

Zebrafish (*Danio rerio*) possess many features that make for an ideal model of embryonic and adult hematopoiesis (Boatman *et al*, 2013; de Jong & Zon, 2005; Paik & Zon, 2010; Stachura & Traver, 2011; Traver *et al*, 2004), including high conservation with the human hematopoietic system, external fertilization, high fecundity, rapid development, and embryonic transparency. A whole genome duplication event in teleosts following evolutionary separation from the mammalian lineage resulted in two zebrafish granulin orthologues: granulin a, *grna*, and granulin b, *grnb* (Cadieux *et al*, 2005). We have taken advantage of the tissue-specific segregation of the zebrafish granulin paralogues to assess the functional role of Granulin in hematopoiesis without perturbing other tissues. We demonstrate that whereas *grnb* is widely expressed in most tissues, *grna* expression is restricted to myeloid cells. Loss of function experiments using specific morpholinos or *grna* and *grnb* null mutants, in combination with *in vivo* imaging and intracellular flow cytometry, demonstrate that *grna*, but not *grnb*, deficiency leads to loss of neutrophils and macrophages due to failure in differentiation from myeloid progenitors during embryonic development. In addition, examination of adult *grna* mutant zebrafish show failure in neutrophil differentiation in the adult hematopoietic system. Mechanistically, we have performed Cleavage Under Targets and Release Using Nuclease (CUT & RUN) and show that the master transcription factor of myeloid differentiation Pu.1 directly binds granulin enhancers, triggering its expression. Moreover, our studies demonstrate that Irf8, one of the main transcription factors regulating macrophage development, also acts upstream of granulin. The regulation of mammalian *GRN* by PU.1 and IRF8 was confirmed using empiric-based databases, highlighting functional conservation between species in myeloid differentiation. Our functional studies *in vivo* demonstrate that loss of *grna* leads to defective recruitment of myeloid cells to wounds due to a lack of mature macrophages and neutrophils. This results in aberrant collagen deposition within the scar and therefore defective wound healing. Altogether, using our zebrafish model of granulin deficiency, we have discovered that granulin is an essential regulator of myeloid cell differentiation. This study therefore opens a new field of investigation that will help shed light on the pleiotropic functions of this enigmatic protein and facilitate its use as a therapeutic target.

## Results

### grna expression is restricted to myeloid cells in the zebrafish embryo

Despite mammalian granulin mRNA being among the most abundant transcripts in human macrophages and other myeloid cell lineages (Ong *et al*., 2006), functional studies on its potential roles *in vivo* have not been reported. This prompted us to identify an animal model of granulin deficiency amenable to determine the role of granulin during myelopoiesis. Previous work has suggested that whereas *grnb* is ubiquitously expressed throughout the zebrafish embryo, *grna* is restricted to the sites of embryonic hematopoiesis (Cadieux *et al*., 2005).

We aimed to further determine when and where *grna* and its paralogue *grnb* were expressed during zebrafish development. First, we performed quantitative PCR (qPCR) in zebrafish embryos from 2-72 hpf. As shown in Figure 1A-B, both *grna* and *grnb* transcripts were maternally transferred, since they were detected at 2 hpf, with *grna* expression 100-200 fold less abundant than *grnb* (notice different Y axis scales). Zygotic *grna* and *grnb* transcripts (detected from 9hpf) were 10 times less abundant than maternally transferred *grna* and *grnb*, but were detected throughout all embryonic stages evaluated, with a peak in expression at 72 hpf (Figure 1A-B).

**Figure 1:**
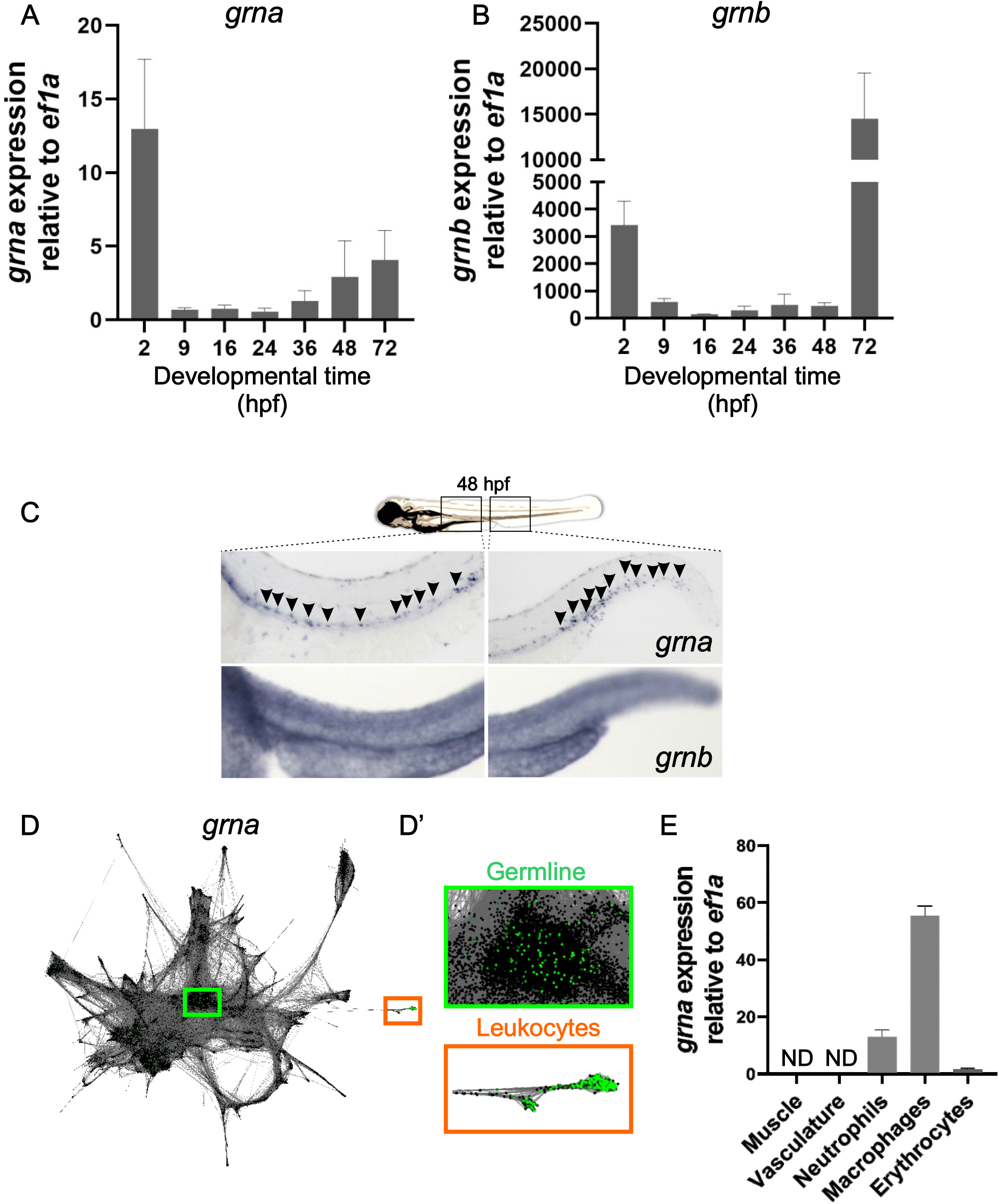
*grna* expression is restricted to the myeloid cell lineage during embryo development. Expression of *grna* **(A)** and *grnb* **(B)** during zebrafish embryonic and larval development. The mRNA levels were determined by real-time quantitative PCR in 10-30 pooled larvae at the indicated times. The gene expression is normalized against *ef1a*, each bar represents the mean ± S.E. from two independent experiments. **(C)** Expression of *grna* (*upper panel*) and *grnb* (*lower panel*) by WISH at 48hpf. Black arrowheads denote *grna* expression by distinct individual cells. Note that the *grnb* expression pattern is ubiquitous. Anterior is to the left, dorsal to the top. **(D-D’)** scRNA-seq graph showing *grna* expression (green dots) using the online tool SPRING by Wagner and colleagues (Wagner *et al*., 2018). Dots represent single-cells from 4 hpf (center) to 24 hpf (periphery) zebrafish embryos. The cells that expressed *grna* (green dots) are magnified in D’. Notice that *grna* expression is restricted to germline cells (green box) and leukocytes (orange box). **(E)** Muscle (*Myf5:eGFP*^+^), vasculature (*Flk:mcherry*^+^; *Gata1:Dsred*^-^), neutrophils (*Mpx:eGFP*^+^), macrophages (*Mpeg1:eGFP*^+^), and erythrocytes (*Globin:eGFP*^+^) cells from dissected embryos were purified by FACS, and qPCR for *grna* performed. Levels of *grna* transcripts along x-axis are shown relative to the housekeeping gene *ef1a*. Bars represent means ± S.E.M. of two to three independent samples. ND, not detected; hpf, hours postfertilization.

To evaluate where in the embryo *grna* and *grnb* were expressed, we generated full-length antisense and sense probes for each granulin paralogue and performed whole mount *in situ* hybridization (WISH). *grna* and *grnb* were expressed by all cells in the early zygote (2-10 hpf) (Supplementary Figure 1). However, *grna* transcripts were restricted to the embryonic hematopoietic areas in the zebrafish at later developmental stages (48 hpf), including the dorsal aorta (DA) region, and caudal hematopoietic tissue (CHT) (Figure 1C, left and right upper panels respectively, black arrowheads). In contrast, *grnb* was detected throughout the embryo (Figure 1C, lower panels). To confirm the cellular origin of *grna* expression throughout embryonic development, we utilized SPRING, a tool for interactive exploration of single-cell data from zebrafish embryos (Wagner *et al*, 2018). Figure 1D-D’ shows a single-cell graph of all individual cells across all time points: from 4 hpf, center; to 24 hpf, outer dots. Cells that expressed *grna* (green dots) are shown in Figure 1D, and magnified view in 1D’. *grna* expression was therefore restricted to germline cells (green box) in the early zygote and leukocytes (orange box) at 24 hpf, further validating our WISH results. In contrast, *grnb* was found to be expressed at low levels in all tissues (data not shown).

We next validated the restricted tissue specificity of *grna* in the late embryo (36-48 hpf) by qPCR of FACS-sorted hematopoietic and non-hematopoietic cells. *grna* was highly expressed by myeloid cells and absent in non-hematopoietic cells such as muscle and endothelial cells (Figure 1E). Among the myeloid cells expressing *grna*, macrophages expressed five times more *grna* transcripts than neutrophils, and 50 times more than developing erythrocytes (Figure 2E).

**Figure 2:**
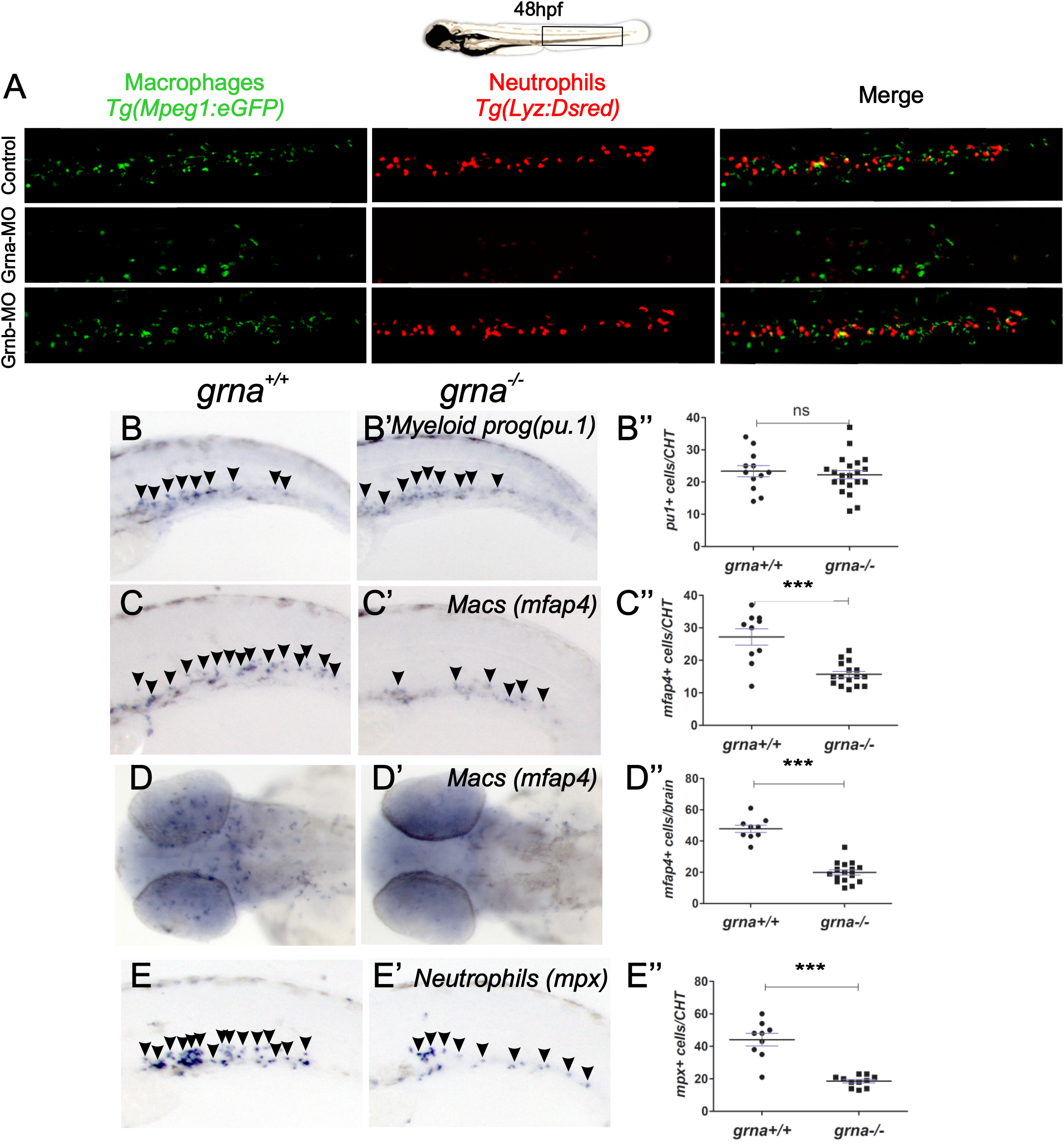
Absence of *grna* leads to decreased myeloid differentiation during embryo development. **(A)** Representative fluorescence images of the tails of 48 hpf *Mpeg1:eGFP; Lyz:dsred* double transgenic embryos injected with Grna mismatch control, Grna or Grnb MOs. **(B, B’-F, F’)** WISH for the myeloid progenitor (*pu.1*), macrophage (*mfap4*) and neutrophilic (*mpx*) markers in *grna*^-/-^ and *grna*^+/+^ control embryos at 48 hpf **(B, B’’-E, E’’)** or 5 dpf **(F-F’’)**. Black arrowheads depict cells expressing the indicated transcripts. **(B’’, C’’, D’’, E’’, F’’)** Enumeration of *pu1^+^, mfap4^+^*, *mpx^+^* and *apoeb^+^* expressing cells shown in (B,B’-F, F’). Each dot represents the number of positive cells in the photographed area in each embryo. Bars represent mean ± S.E.M. Ns, not significant. *p < 0.05, and ***p < 0.001.

Altogether, these results demonstrate that in zebrafish, *grna* expression is restricted to the myeloid cell lineage from 24 hpf, while *grnb* is ubiquitously expressed throughout the embryo.

### Grna is required for proper myeloid development

Due to the restricted expression of *grna* to developing myeloid cells in the embryo, we hypothesized that *grna* could be essential for proper myeloid differentiation. To test our hypothesis, we first performed loss-of-function experiments for *grna* and *grnb* utilizing specific antisense morpholinos (MOs). In the zebrafish embryo, myeloid progenitors, macrophages, and neutrophils can be visualized as discrete cells by expression of *pu.1, mfap4*, and *mpx*, respectively, using WISH or the zebrafish transgenes *Tg(pU1:Gal4; UAS:eGFP), Tg(Mpeg1:eGFP)*, and *Tg(Lyz:Dsred)*, respectively. Grna knock-down utilizing a previously validated MO (Li *et al*, 2010) (Supplementary Figure 2A) significantly reduced *in vivo* macrophage (*mfap4*) and neutrophil (*mpx*) numbers at 48 hpf compared to *grna* mismatch control MO injected embryos (Figure 2A upper and middle panels). To investigate if the absence of Grnb also led to myeloid defects, we generated and validated by RT-PCR a specific *grnb* splice-blocking MO (Supplementary Figure 2C-D). Although some WT *grnb* transcripts were observed after Grnb-MO injection (Supplementary Figure 2C, lower band), this amount was significantly reduced compared to WT control embryos. No significant changes in neutrophil or macrophage numbers were observed after *grnb* knock-down (Figure 2A, bottom panel). We further confirmed the loss of macrophages and neutrophils after *grna* knockdown by using an additional *grna* MO (grna-MO2) (Li *et al*., 2010) and performing flow cytometry in pooled *Mpeg1.1:eGFP* 36hpf embryos (Supplementary Figure 2D-E). As expected, macrophage numbers were significantly reduced in embryos injected with either Grna MO1 or MO2 morpholinos compared to control Grna mismatch MO injected embryos.

To verify that the loss of macrophages and neutrophils was not due to potential off-target effects caused by the *grna* morpholinos, we generated *grna* null mutants (Solchenberger *et al*, 2015) and performed WISH for *pu.1, mfap4* and *mpx* at 48hpf. While myeloid progenitor (*pu.1*+) numbers were similar (Figures 2B-B”), macrophage and neutrophil numbers were significantly decreased in *grna*^-/-^ compared to *grna*^+/+^ control embryos (Figures 2C-E”). We next performed intracellular flow for Mfap4 to further validate the decrease in macrophages. As expected, the number of Mfap4^+^ cells was significantly decreased (5-fold) in *grna*^-/-^ at 48 hpf (Supplementary Figure 2F-G). To ensure that the myeloid defects observed in the absence of *grna* were not due to a decreased proliferation of differentiated myeloid cells, we performed whole-mount immunohistochemistry (WIHC) for phospho-histoneH3 (P-H3) in 30 hpf *Mpx:eGFP* transgenic animals and found no colocalization of *Mpx:eGFP* with P-H3 in *grna*^+/+^ embryos, indicating that neutrophils do not proliferate during early embryonic development (Supplementary Figure 2H). Altogether, these experiments demonstrate that *grna*, but not *grnb*, is essential for the differentiation of myeloid progenitors into macrophages and neutrophils.

### Absence of grna leads to long lasting myelopoiesis defects

We next sought to investigate if *grna* was also essential for adult myelopoiesis or, in contrast, if these defects were specific to embryonic development. First, we investigated if *grna* expression was restricted to the myeloid lineage in adult vertebrates. To address this question, we took advantage of the interactive scRNAseq data published by Tang and colleagues (Tang *et al*, 2017) from adult wildtype zebrafish kidneys, the hematopoietic zebrafish analogue to mammalian bone marrow. *grna* expression was restricted to the cell clusters defined as macrophages (yellow circles) and myeloid cells (green circle) (Figure 3A and Supplementary Figure 3A). In contrast, *grnb* transcripts were present at low levels in most hematopoietic cells, but highly enriched in multicilliated cells and vascular endothelium (Sup Figure 3A), which are non-hematopoietic cells within the kidney. We next wanted to address if, similar to the zebrafish orthologue *grna*, mammalian granulin expression was also up-regulated in myeloid cells. First, we utilized *Gene Expression Commons* (Seita *et al*, 2012) to query the dynamic-range of granulin within microarrays of the mouse (*Mus musculus*) hematopoetic system (https://gexc.riken.jp/models/3/genes/Grn?q=Grn). As shown in Figure 3B, *Grn* expression was active in hematopoietic stem cells (HSCs), up-regulated in granulocyte/macrophage progenitors (GMPs) and reached its highest expression in granulocytes (Gra) and monocytes (Mono). In contrast, cells of the megakaryocyte/erythrocyte lineage, including megakaryocyte/erythrocyte progenitors (MEPs), Megakaryocyte progenitors (Mkps), and Colony Forming Unit-Erythroid (PCFU-e) drastically down-regulated *Grn* expression. Second, to confirm these results at the single cell level, we utilized the hematopoietic single cell interactive gene viewer based on mouse sorted hematopoietic cells published by Olsson and colleagues (Olsson *et al*, 2016) and found that murine *GRN* is highly expressed in myeloid cells including monocytes, granulocytes, and myelocytes, mimicking the expression of the master myeloid transcription factor PU.1 (Supplementary Figure 3B, black squares). Both *Grn* and *Pu.1* were down-regulated in cells of the erythroid lineage (Supplementary Figure 3B, purple squares). Altogether, these data show conserved expression for the mammalian granulin and the zebrafish orthologue *grna* in hematopoietic lineages.

**Figure 3:**
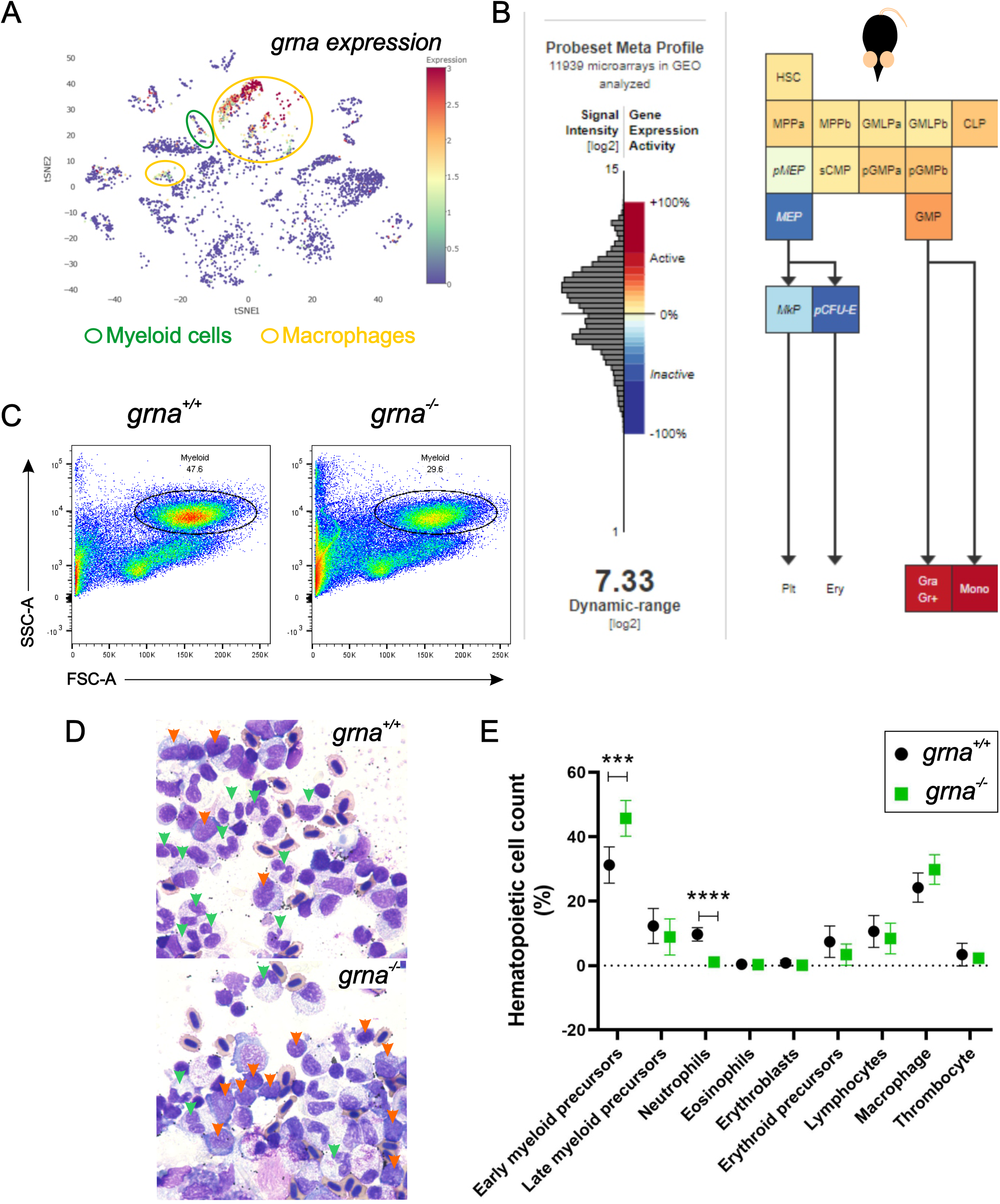
Granulin expression is up-regulated in vertebrate myeloid cells and is essential for myeloid cell differentiation during adult hematopoiesis. **(A)** tSNE analysis showing *grna* expression levels (red is high, orange and yellow are medium expression; blue is absent) of single cells sequenced from wildtype zebrafish kidney marrow using the online visualizer “Single Cell inDrops RNA-Seq Visualization of Adult Zebrafish Whole Kidney Marrow” (https://molpath.shinyapps.io/zebrafishblood/#pltly) (Tang *et al*., 2017). The main tSNE clusters identified expressing *grna* are denoted by open circles. Yellow open circles represent clusters defined as *“macrophages”*. Open green circles are *grna* expressing clusters whose cells were identified as *“myeloidc ells”*. **(B)** Mouse hematopoietic model showing the dynamic expression of Grn derived from microarray data (Affymetrix Mouse Genome 430 2.0 Array). Notice that lymphocyte differentiation beyond CLP is not shown here. **(C)** Representative flow cytometric light scatter profile showing the different hematopoietic populations present in *grna*^+/+^ (left) and *grna*^-/-^ (right) kidney marrow. **(D)** Representative pictures from *grna*^+/+^ and *grna*^-/-^ whole kidney marrows cytospins stained with Wright-Giemsa showing increased early myeloid precursors (orange arrowheads) and decreased mature neutrophils (green arrowheads) in the absence of *grna* (bottom panel) compared to *grna*^-/-^ control siblings (upper panel). **(E)** Manual quantification of kidney marrow hematopoietic cells in *grna*^-/-^ (green squares, n=5) compared to control *grna*^+/+^ (black dots, n=5) from two independent experiments. Horizontal lines and error bars indicate means ± SEM (***p<0.001, ****p<0.0001). HSC, Hematopoietic Stem Cell; MPP, Hematopoietic multipotential progenitors; GMLP, granulocyte–monocyte–lymphoid progenitor; CLP, common lymphoid progenitor; pMEP, pre of megakaryocyte-erythroid progenitor; MEP, megakaryocyte-erythroid progenitor; MkP, Megakaryocyte progenitor; PCFU-e, Colony Forming Unit-Erythroid; Plt, platelets; Ery, Erythrocytes; sCMP, strict common myeloid progenitor; pGMP, pre-granulocyte/macrophage; GMP, granulocyte/macrophage progenitors; Gra Gr+, granulocytes; Mono, monocytes.

To assess the effect of *grna* loss on myeloid cell differentiation in the adult vertebrate, we collected whole kidney marrow cells from *grna*^-/-^ and control *grna*^+/+^ siblings and performed flow cytometry to quantify myeloid cell numbers using the forward and side scatters as we described previously (Traver *et al*, 2003). The percentage of myeloid cells within the kidney marrow of *grna*^-/-^ was significantly decreased as compared to *grna*^+/+^ control kidneys (Figure 3C and Supplementary Figure 3C). We next cytospun *grna*^-/-^ and *grna*^+/+^ control sibling kidney cell suspensions and subsequently performed Wright-Giemsa staining to examine hematopoietic counts and morphology. Pathologist examination revealed that *grna*^-/-^ had increased early myeloid precursors (Figure 3D, orange arrowheads) with decreased differentiation into mature neutrophils (green arrowheads) based on 200 non-erythroid nucleated differential cell counts (Figure 3D-E). Importantly, there were no differences in body size between *grna*^+/+^ and *grna*^-/-^ sibling fish as measured by length from mouth to caudal fin (Supplementary Figure 3D). Taken together, these data indicate that *grna* is essential to drive myeloid cell differentiation from early myeloid precursors in adult zebrafish.

### *grna* inhibits erythroid differentiation

Next, we performed bulk RNA-seq in kidney marrow from adult *grna*^-/-^ and *grna*^+/+^ control siblings to identify genes whose expression was dysregulated in the absence of *grna*. We found 154 genes significantly down-regulated in the *grna* mutants (Table S2), and 116 genes significantly up-regulated (Figure 4A-C) (Table S3). As expected, we found important myelopoiesis-related genes down-regulated in *grna* mutants, including *cebpa, rgs2, apoeb* and *abca1b* (Figure 4B-C). Interestingly, when we looked at the cell types that expressed these down-regulated genes using the online scRNA-seq viewer from zebrafish kidney marrow (Tang *et al*., 2017), we found that almost half of them (64 out of 154) (Table S4) were restricted to myeloid cell subpopulations (Sup Figure 4), supporting a role for *grna* in myeloid cell differentiation. In addition, many genes down-regulated in the *grna* mutants have been shown to participate in inflammation and immune response, including *i113ral, il4r.1, irf2a, itgb3a, nfkbie, cp, tlr5b, cxcl12a, hbegfa, mlx* and *atg16l1* (Figure 4B-C and Sup Fig 4). Altogether, this data further confirms that *grna* is essential for proper myeloid cell differentiation.

**Figure 4:**
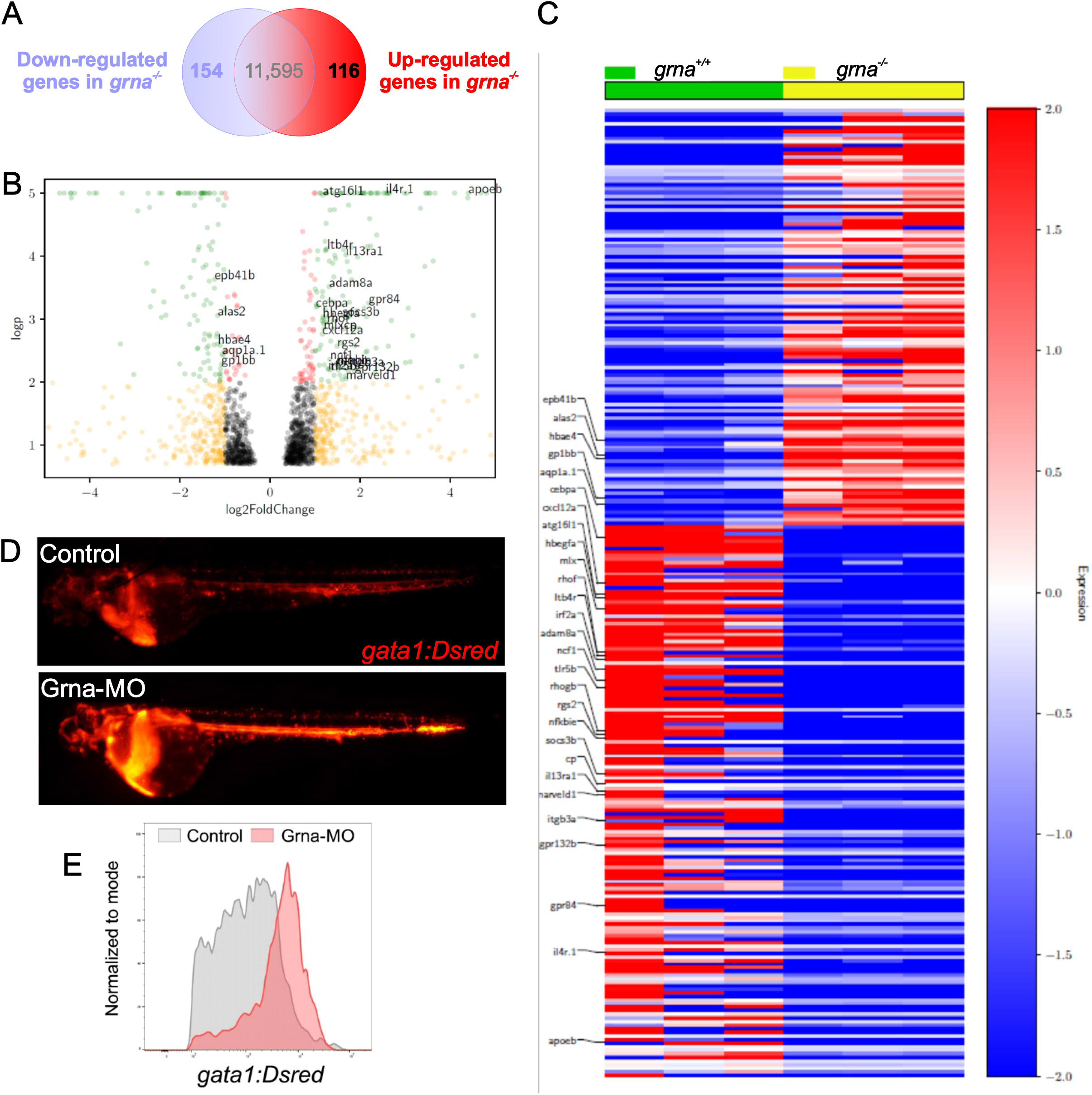
Grna inhibits erythroid differentiation. **(A)** RNA-seq analysis from *grna*^-/-^ and *grna*^+/+^ adult zebrafish kidney marrows reveals 154 down-regulated and 116 up-regulated genes in *grna*^-/-^ versus *grna*^+/+^ control. **(B)** Volcano plot obtained from DESeq2 analysis of *grna*^-/-^ and *grna*^-/-^ kidney marrows. **(C)** Heat map of the enriched and depleted transcripts in kidney marrows from *grna*^-/-^ versus *grna*^-/-^ adult fish. Color-coding is based on rlog-transformed read count values. **(D)** Fluorescence images of *gata1a:Dsred* embryos injected with Grna morpholino (Grna-MO) (upper panel) and a five-base Grna mismatch control (bottom panel), showing increased *gata1a:Dsred* fluorescence in the absence of Grna. **(E)** Histogram of three-pooled *gata1a:Dsred* embryos injected with Grna-MO (red) or mismatch Grna-MO control (grey).

Unexpectedly, among the up-regulated genes in *grna*^-/-^, we found genes described to be important for red blood cell development (*alas2*), erythrocyte shape (*epb41b*), hemoglobin transport (*hbae4*) as well as platelet adhesion (*gp1bb*) (Figure 4B-C). Surprisingly, the expression of 27 out of 116 upregulated genes was found to be restricted to erythrocytes and, to a lesser extent, platelets (Sup Figure 4 and Table S5). We then hypothesized that *grna* might inhibit erythroid differentiation. To address this, we injected Grna-MO and Grna mismatch control MO into *gata1a:DsRed* reporter zebrafish embryos. Gata1 is a master transcription factor that drives erythroid cell differentiation. We found a significant increase in Dsred expression in Grna morphants compared to control siblings (Figure 4D), suggesting that *grna* inhibits *gata1a*. We next quantified this up-regulation using flow cytometry and found a 10-fold increase in DsRed intensity in the absence of *grna* (Figure 4E), indicating that the *gata1a* promoter is 10 times less active *in vivo* in the absence of Grna. Altogether, these results show that Granulin inhibits erythroid differentiation and facilitates myeloid cell development *in vivo*.

### Pu.1 and Irf8 control grna expression in zebrafish, and this regulatory mechanism is conserved for mammalian GRN

Pu.1, encoded by the *Sfpi1* (*Spi-1*) gene, is a master transcription factor that leads to myeloid cell specification. Pu.1 contains various Ets domains that recognize the DNA sequence harboring the core GGAA motif (Burda *et al*, 2010) (Figure 5D). Since *pu.1* expression was unaltered in the absence of *grna* (Figures 2B-B”), we hypothesized that *grna* acted downstream of Pu.1 for myeloid cell specification. First, to address if *grna* and *pu.1* transcripts were co-expressed by the same cells, we performed double fluorescence *in situ* hybridization (FISH) for *grna* and *pu.1* in 48 hpf embryos. Figure 5A shows co-localization of *grna* and *pu.1* at a single cell resolution using confocal microscopy. To demonstrate that Pu.1 genetically acts upstream of *grna*, we utilized a specific MO against Pu. 1 (Rhodes *et al*, 2005) and performed WISH for *grna* and the neutrophilic marker *mpx*. As shown in Figure 5B (left and middle panels), Pu.1 knockdown completely abolished *grna* expression, as well as decreased the neutrophilic marker *mpx* as previously reported (Espin-Palazon *et al*., 2014). To control for the efficiency of the Pu.1-MO, we injected *Mpeg1:eGFP* embryos and found that macrophages were completed ablated in the absence of Pu.1 as expected (Supplementary Figure 5A). To address if Pu.1 directly bound *grna* enhancers to promote its transcription, we utilized the motif comparison tool *Tomtom* (http://meme-suite.org/tools/tomtom) (Bailey *et al*, 2009) to identify putative Pu.1 binding sites in *grna* regulatory sequences. Using the PU.1 matrix ID: MA0080.5 http://jaspar.genereg.net/matrix/MA0080.5/ from *Homo sapiens* derived from ChIP-seq (Chromatin immunoprecipitation followed by sequencing) data (Ray-Gallet *et al*, 1995) (Figure 5D), we found several putative binding sites (BS) in the non-coding regions flanking the *grna* gene (Figure 5C). These included three predicted BS in *grna* intron 3-4 (BS1 at +5655, BS6 at +4485); and three in the *grna* 5’ enhancer region (BS2 at −6935, BS5 at −7843, BS3 at −14045) (Figure 5C). We then dissected zebrafish kidney marrow to perform Cleavage Under Targets and Release Using Nuclease (CUT&RUN) with a specific antibody for zebrafish Pu.1 followed by qPCR to amplify each predicted pU.1 BS. We chose CUT&RUN as a chromatin profiling strategy since it is robust and has extremely low background signal (Skene & Henikoff, 2017). While putative Pu.1 BSs 5, and 6 showed no enrichment compared to control isotype IgG antibody, Pu.1 BS1, BS2 and BS3 showed two-to eight-fold enrichment (Figure 5E), demonstrating that Pu.1 directly binds *grna* regulatory sequences. To gain further insight into the *grna* regulatory gene network, we knocked down Irf8, an essential transcription factor required for macrophage specification. It has been demonstrated that Pu.1 acts upstream of Irf8, and that both transcription factors cooperate to regulate granulocyte-macrophage fate decisions in myeloid progenitors and maturation of macrophage precursors in both zebrafish and mice (Kurotaki *et al*, 2018; Li *et al*, 2011; Shiau *et al*, 2015).

**Figure 5:**
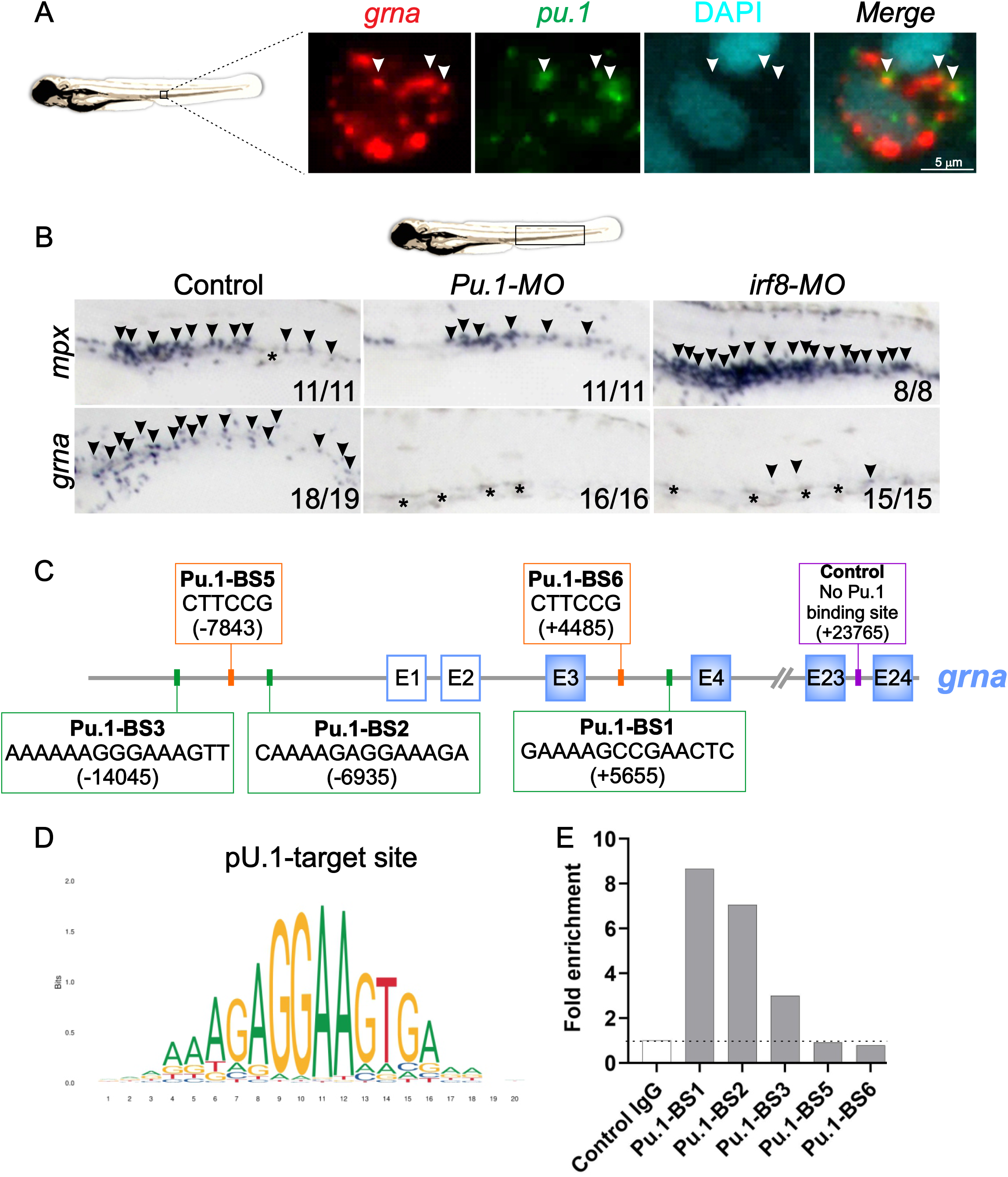
Transcriptional factor network that controls granulin expression. **(A)** Double fluorescence *in situ* hybridization for *grna* (red) and *pu.1* (green) shows colocalization of both transcripts. Nuclei are stained with DAPI (blue). Pictures were taken by confocal microscopy from the tail region of 48 hpf zebrafish embryos. Each image is a 1 μm z slice. **(B)** WISH for the neutrophilic marker *mpx* (upper panels, arrowheads) or *grna* (lower panels, arrowheads) after morpholino knockdown of Pu.1 (middle panels) or Irf8 (right panels) compared to Standard morpholino control (left panels) at 48 hpf. Pu.1 or Irf8 knockdown abolished *grna* expression. Asterisks denote natural pigmentation occurring in the tail of zebrafish embryos. **(C)** Schematic representation of the *grna* gene and its 5’ enhancer locus denoting six putative Pu.1 binding sites (BSs). Pu.1 BS1-3 (green squares) were found searching the human pU.1-target site nucleotide matrix represented in **(D)** using the motif comparison tool Tomtom. Pu.1 BS5-6 (orange squares) were found searching the pU.1-target site nucleotide matrix available in ConSite (http://consite.genereg.net/cgi-bin/consite). BSs positions are denoted with numbers in between brackets. Positive “+” numbering starts with +1 at the A of the *grna* ATG translation initiation (start) codon. Nucleotides upstream (5’) of the *grna* ATG-translation initiation codon (start) are marked with a “-” (minus). E, exon. White squares with blue line indicate *grna* exons from the untranslated region. Blue squares represent *grna* coding exons. Control primers to amplify 71 base pairs of a locus within the *grna* gene with no predicted Pu.1 BSs for CUT&RUN normalization are indicated with a purple square. **(E)** Representative CUT&RUN experiment performed in fresh zebrafish kidney marrows from adult AB* using a Pu.1 or control IgG antibody. Fold enrichment of Pu.1-associated DNA fragments were identified by qPCR using primers flanking the BSs denoted in **(C)**. To calculate the fold enrichment, qPCR results for each BS were normalized against spike-in DNA as described (Meers *et al*, 2019) and control primers that amplify a locus of the *grna* gene lacking predicted Pu.1 BSs. This experiment was performed three independent times with similar enrichments. The primers used are shown in Table S1.

Irf8 depletion by utilizing a specific Irf8 morpholino (Li *et al*., 2011) led to increased *mpx* expression (Figure 5B, right top panel) and loss of macrophages (Supplementary Figure 5A) as previously reported (Espin-Palazon *et al*., 2014). In addition, Irf8 knockdown resulted in a significant decrease in *grna* expression (Figure 5B, right bottom panel). Using Gene Expression Commons (Seita *et al*., 2012) to query the dynamic-range of *Irf8* expression within microarrays of the mouse (*Mus musculus*) hematopoietic system, we found that Irf8 is up-regulated in myeloid precursors (Supplementary Figure 5B). Together, these results indicate that Irf8 acts genetically upstream of *grna* to trigger *grna* expression within myeloid precursors.

We next wanted to investigate if the transcriptional network that regulated *grna* in zebrafish myeloid cells was also conserved in mammals. We queried if PU.1 bound the human *GRN* promoter by using the regulatory feature of *ensembl.org* (Zerbino *et al*, 2016). This database uses a variety of published genome-wide assays, such as ChIP-seq data on chromatin structure, epigenomic, and transcription factor binding to identify peaks in the genome that correspond to binding of a TF. We found that in human cells, PU.1 bound to the first intron of the human *GRN* gene (Supplementary Figure 5C). Next, we utilized ChIP Enrichment Analysis (ChEA) (Lachmann *et al*, 2010) to search for other TFs binding the mammalian *GRN* regulatory regions. ChEA utilizes transcription factor ChIP-seq studies extracted from supporting materials of publications to identify peaks at the promoter of a queried gene. We found that the myeloid TFs IRF8 and CEBPB bound the *GRN* promoter in mouse with significant P-values<0.05 (Supplementary Figure 5D). In addition, hematopoietic and inflammatory TFs including RUNX1, Growth Factor Independent 1 Transcriptional Repressor (GF1), Nuclear Factor, Erythroid 2 Like 2 (NFE2L2) and RELA Proto-Oncogene, NF-KB Subunit (RELA) were also detected to bind mammalian *GRN*, although with lower significance (Supplementary Figure 5D). We next used Harmonizome (Rouillard *et al*, 2016) in combination to ChEA to identify genes that were co-expressed with *GRN*. As expected, mammalian *PU.1* and *IRF8* were co-expressed with *GRN* (Supplementary Figure 5E), as well as myeloid specific genes such as *MPEG, CD68*, and *TREM2* with a Pearson Correlation >0.6 (Supplementary Figure 5F).

Taken together, these data demonstrate that in zebrafish, Pu.1 and Irf8 positively control *grna* expression in myeloid cells, and that this regulatory mechanism is conserved for human and mouse granulin.

### grna is required for emergency myelopoiesis and macrophage recruitment to the wound

Emergency myelopoiesis is the proliferation and differentiation of hematopoietic progenitor cells towards the myeloid lineage as a result of increased demand following injury or infection (Mitroulis *et al*, 2018). We have shown that myeloid progenitors are present in the absence of *grna* (Figures 2B-B’ and 4F-G). Because *grna* is required for normal myelopoiesis, we hypothesized that the absence of *grna* would lead to an aberrant emergency myelopoiesis response. To investigate if myeloid progenitors could respond to emergency myelopoiesis, transgenic *Mpeg1:eGFP* embryos were injected with Grna or Grna control mismatch MOs and caudal tails were resected at 48 hpf to trigger an emergency myelopoiesis response. Fluorescent images were taken from 1 hour-post wounding (hpw) to 32 hpw (80 hpf), and macrophage numbers were quantified in the tail (CHT and wound) (Figure 6A). While macrophage numbers increased in all individuals injected with Grna mismatch control morpholino at 9 hpw, Grna morphants failed to generate more macrophages (Figure 6B), suggesting that Grna is required for emergency myelopoiesis. Although the absence of *grna* led to a remarkable decrease in macrophage number, a small percentage of macrophages was still present in *grna* mutants and morphants (Figure 2). Therefore, we sought to determine if these macrophages were able to respond and recruit to the injury site using the same experimental approach described above (Figure 6A). As expected, the number of macrophages recruited to the wound was significantly decreased in the absence of Grna (Figure 6D). This could be due to the reduced numbers of macrophages in the Grna morphants, but the percentage of recruited macrophages with respect the total macrophage number per individual could be normal. To address this, we quantified the number of macrophages recruited to the wound and normalized it to the total number of macrophages in the tail (CHT and wound) for each individual. The percentage of macrophages that were recruited to the wound with respect to the total macrophage numbers was significantly reduced in the absence of Grna (Figure 6C) compared to control Grna mismatch morpholino injected siblings. Taken together, these results indicate that Grna is essential for emergency myelopoiesis and that the macrophages produced in the absence of Grna have functional abnormalities, not being able to properly respond to tissue injury.

**Figure 6:**
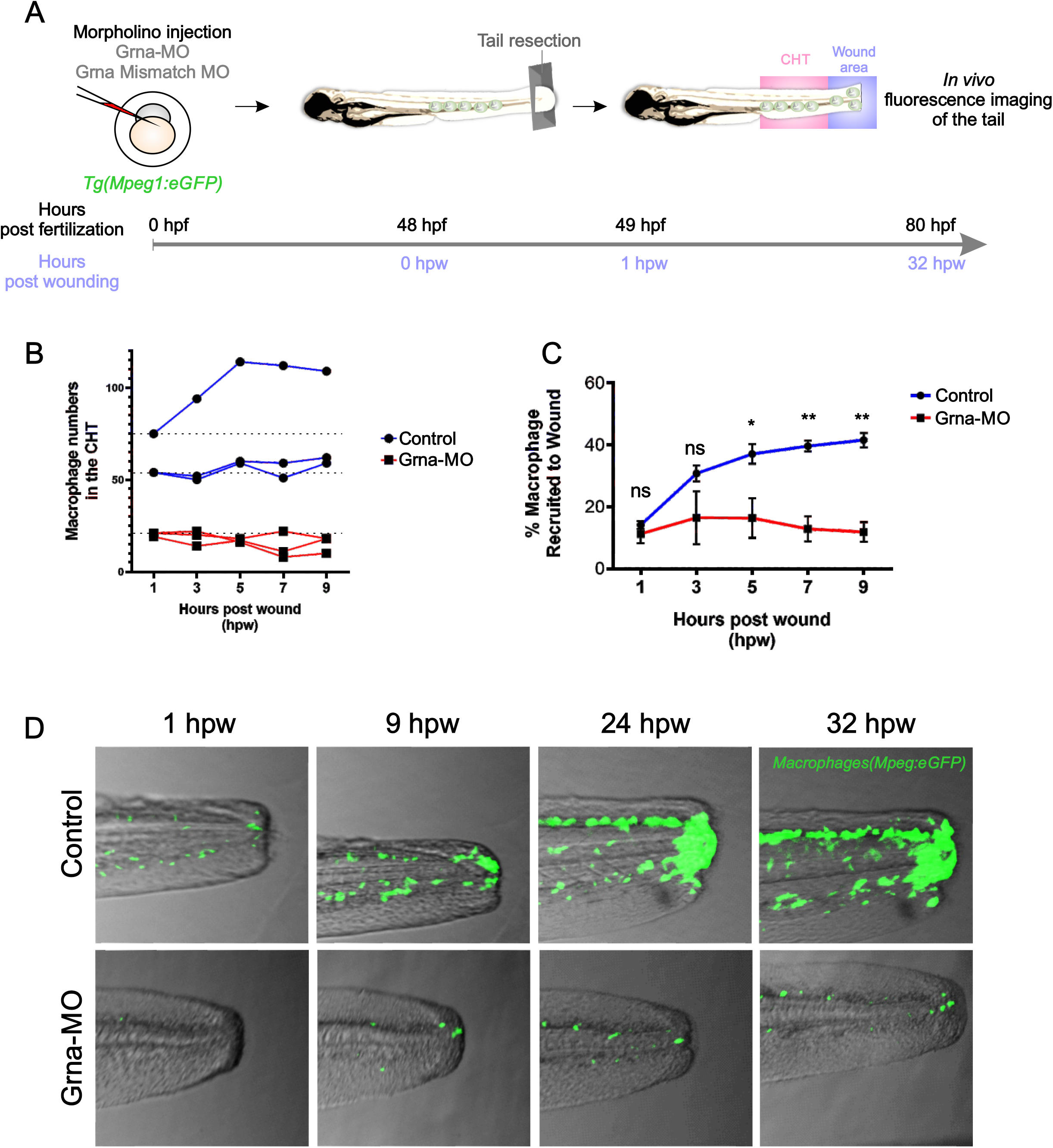
Macrophages respond abnormally to injury in the absence of Grna. **(A)** Experimental workflow. Transgenic *Mpeg1:eGFP* one-cell stage embryos were injected with either GrnaMO or Grna mismatch MO. At 48 hpf, caudal tails were resected immediately after the end of the notochord. Fluorescence imaging of the tail (CHT, were the majority of neutrophils reside at this developmental time, and wound area) was taken every two hours from 1 hpw to 32 hpw, and the number of neutrophils quantified manually. **(B)** Neutrophil numbers in the CHT from individual *Mpeg1:eGFP* transgenic animals at 1, 3, 5, 7 and 9 hpw following depletion of Grna compared to Grna mismatch control morphants. **(C)** Percentage of *Mpeg1:eGFP^+^* macrophages recruited to the wound region normalized to the total macrophage number in the tail (CHT and wound) in GrnaMO (red line, n=3) and Grna mismatch MO (blue line, n=3) injected embryos at indicated time points. Circle and square dots indicate means, and error bars SEM (ns, not significant; *p<0.05, **p<0.001). **(D)** Representative fluorescence images of tail fins from *Mpeg1:eGFP* Grna- or Grna mismatch control morphant siblings at the indicated times.

### Wound healing is impaired in grna mutants

It has been recently demonstrated that macrophages are one of the main contributors to tissue repair in both mammals and zebrafish (Li *et al*, 2012; Minutti *et al*, 2017; Miskolci *et al*, 2019; Morales & Allende, 2019; Nguyen-Chi *et al*, 2017; Simoes *et al*, 2020). They contribute directly and indirectly by ingesting apoptotic neutrophils, triggering the resolution of inflammation, breaking down the injured matrix, re-epithelialization, vascularization of the wound bed, releasing growth factors, induction of extracellular matrix deposition (Minutti *et al*., 2017), and even depositing collagen themselves as shown recently (Simoes *et al*., 2020). In zebrafish, specific ablation of macrophages severely impedes inflammatory resolution and tissue regeneration (Li *et al*., 2012). Moreover, it is known that Granulin facilitates wound healing by increasing macrophage, fibroblast, and blood vessel number in the injured tissue (He *et al*., 2003). Based on these data, we hypothesized that wound healing would be impaired in *grna*^-/-^ embryos due to the decreased and abnormal production of macrophages, and that the functions attributed to Granulin during wound healing were the consequence of the myelopoietic defects described herein (Figures 2–4). To test our hypothesis, we performed tail fin resection at 48 hpf in *grna*^-/-^ and *grna*^+/+^ control embryos and subsequently imaged regenerated tissue over three days (72hpw) (Figure 7A). Tissue regeneration was deeply impaired in *grna*^-/-^ larvae as shown in Figure 7B. First, the area of tissue regenerated was significantly diminished in *grna*^-/-^ compared to *grna*^+/+^ at all times evaluated (Figure 7C). While *grna*^+/+^ control larvae completely regenerated the tail fin at 72 hpw, *grna*^-/-^ failed to do so (Figure 7B-C). Second, collagen organization in the regenerated area of *grna*^-/-^ was deeply disrupted as shown in Figure 7D. While collagen fibers in *grna*^+/+^ control individuals were perpendicular to the wound edge (Figure 7D, left panel), aligned fibers were disarrayed and less perpendicular to the wound edge in *grna*^-/-^ (Figure 7D, right panel). These observations are consistent with, and extend previous findings that macrophages indirectly and directly support collagen fiber production and organization during wound repair. Altogether, these results link for the first time the role of granulin in wound healing with the myelopoietic defects described here, and validate functional conservation between the mammalian granulin and the zebrafish orthologue *grna*. Overall, these data also reveal the TFs controlling *grna* expression and the proteins that act downstream of Grna to facilitate myeloid cell differentiation and inhibit the erythroid program (Figure 7E).

**Figure 7:**
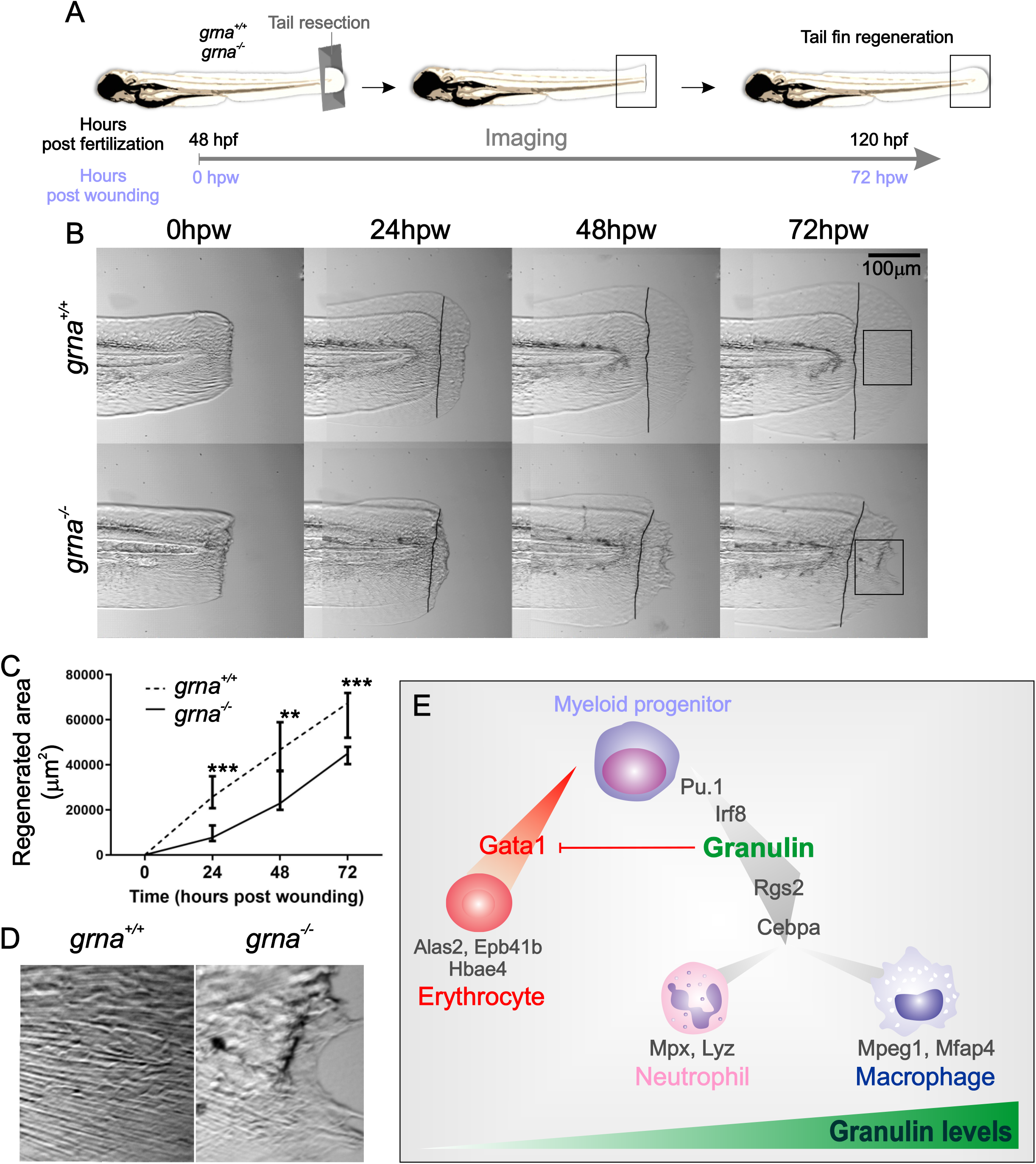
Grna mutants fail to regenerate the tail fin after resection. **(A)** Experimental workflow. *grna*^+/+^ or *grna*^-/-^ 48 hpf embryos were subjected to caudal tail resection immediately after the end of the notochord. Bright field imaging of the wound was taken every 24 hours for three days (72 hpw, equivalent to 120 hpf). **(B)** Representative images of fin tails from *grna*^+/+^ (top panel) or *grna*^-/-^ (bottom panel) larvae at the indicated times. Black lines indicate where the fin tails were resected. **(C)** Quantification of the regenerated tail fin area of *grna*^+/+^ (n=5) and *grna*^-/-^ (n=5) larvae from **(B)**. **p<0.001, ***p<0.0001. **(D)** Magnification of regenerated tail fins from *grna*^+/+^ and *grna*^-/-^ larvae (black rectangles in B). To facilitate collagen fivers visualization, raw pictures were contrasted automatically using the *“contrast enhancement”* tool of CorelDraw version X6. **(E)** Schematic representation of signaling occurring during myeloid cell differentiation. Briefly, Pu.1 and Irf8 positively regulate granulin expression, which in turn controls the expression of *rgs2* and *cebpa* for the differentiation of myeloid progenitors into neutrophils expressing *mpx* and *lyz* or macrophages (*mpeg1* and *mfap4)*. Granulin blocks *gata1* expression, inhibiting erythroid development and the expression of the erythroid related genes *alas2, epb41b* and *hbae4*. Granulin expression levels are indicated in green.

## Discussion

Granulin profoundly impacts wound healing, autoimmune diseases, tumorigenesis, aging, and lysosome biology. In addition, loss of function mutations in the granulin gene are causative for Frontotemporal Lobar Degeneration (FTD) and Ceroid Lipofuscinosis (Petkau & Leavitt, 2014; Ward *et al*, 2017). The tremendous implications that granulin function has in human health, have put this enigmatic protein in the spotlight of many investigations to utilize it as a therapeutic target. Despite its medical importance, why and how granulin can influence these diverse biological processes have remained unclear. Here, we have uncovered a new *in vivo* function for granulin in myeloid cell differentiation. We revealed that loss of granulin causes less macrophages and neutrophils to differentiate with consequences for wound healing and inflammation.

In contrast to mammalian granulin, which is highly expressed by myeloid cells but also present at lower levels in non-hematopoietic tissues, we have demonstrated that the zebrafish granulin orthologue *grna* is restricted to hematopoietic cells, while *grnb* is expressed in most tissues. Our data suggests that *grna* evolved to specifically take on the hematopoietic functions of the mammalian granulin. The tissue-specific segregation of the zebrafish granulin paralogues has therefore allowed an unprecedented manner of assessing granulin function in hematopoiesis without perturbing other tissues. Using our zebrafish model of *grna* deficiency, we have demonstrated that *grna* is essential for neutrophil and macrophage differentiation from myeloid progenitors during normal and emergency myelopoiesis, therefore impacting inflammation and wound repair. Importantly, we show that the regulatory mechanisms of expression of *grna* and mammalian granulin by hematopoietic cells is highly conserved among zebrafish and mammals.

In the absence of *grna*, we persistently observed a profound defect in neutrophil differentiation, while some macrophages still developed. Our data support a functional and transcriptional dysregulation in macrophages in the absence of *grna*, since: (1) Macrophages failed to recruit to the wound; (2) new macrophages failed to generate during emergency myelopoiesis; and (3), many of the down-regulated genes in the *grna*^-/-^ vs. *grna*^+/+^ kidneys are known anti-inflammatory genes, and some completely restricted to macrophages (*abca1b, il4r1*). This aberrant production of inflammatory genes in the absence of *grna* indicates a dysregulated inflammatory response, which is consistent with previous reports in mice lacking *Grn* (Yin *et al*., 2010) and our previous data showing morphological and transcriptional changes in microglial cells, the resident macrophages of the brain, in zebrafish lacking *grna* and *grnb*, indicative of a pro-inflammatory phenotype (Zambusi *et al*, 2020). These observations also support functional conservation between the mammalian granulin and the zebrafish orthologue *grna*.

The transparency of zebrafish larvae allowed us to perform functional assays to visualize the recruitment of macrophages to wounded tissue and its repair. We demonstrate that wound healing is abnormal in individuals lacking *grna*, as previously reported in granulin knockout mice (He *et al*., 2003). However, the restricted *grna* expression within hematopoietic cells in our zebrafish model of *grna* deficiency allowed us to address that the defects in wound healing are caused, at least mostly, by aberrant myeloid cell differentiation. In accordance to our data in zebrafish, it has been reported that the majority of the granulin expressed in the wound derived from myeloid cells (He *et al*., 2003). Since macrophages greatly contribute to tissue repair, it is not surprising that impaired myeloid differentiation leads to aberrant healing of the wounded tissue. Altogether, we have been able to reproduce previously described phenotypes of mice models of granulin deficiency in our *grna* zebrafish mutants and address that the cause of aberrant inflammation and wound healing is due to impaired myelopoiesis.

Our work here also opens the outstanding question of whether the contributions described for granulin in tumorigenesis (Arechavaleta-Velasco *et al*, 2017) are also due to a likely abnormal production of tumor associated neutrophils (TANs), macrophages (TAMs), and antitumor macrophages that constitute the tumor microenvironment, impacting the tumor progression (Pathria *et al*, 2019). In addition, our discovery here paves the way for the understanding of how granulin leads to frontotemporal dementia. In the last years, multiple lines of evidence reveal the impact of microglia in neurological disorders (Bachiller *et al*., 2018; Heneka, 2019). The production of aberrant microglia in the absence of granulin has also been described recently in the adult zebrafish (Zambusi *et al*., 2020) and mice (Bright *et al*, 2019; Lui *et al*, 2016), being the cause rather than the consequence of neurodegeneration. A plausible hypothesis is therefore that microglia-driven neuroinflammation in FTD is caused by the myeloid defects described here in myeloid development. Altogether, our results suggest genetic and functional differences in macrophages produced in the absence of *grna*, which most likely impact the homeostasis of many tissues due to the ubiquitous presence of resident macrophages and the implications that macrophages and neutrophils have in tumor progression, autoinflammatory disease and neurodegenerative disease. More experiments in each of the contexts will be needed to address the exact differences among microglia or tumor associate myeloid cells in the absence of granulin, and our zebrafish model of granulin deficiency is an ideal model to perform these studies *in vivo*.

Among the complex regulatory network of TFs that regulate hematopoiesis, GATA1 and PU.1 are key on regulating the erythroid versus myeloid program by antagonizing each other (Iwasaki & Akashi, 2007; Monteiro *et al*, 2011; Wolff & Humeniuk, 2013; Zhang *et al*, 2000). These TFs are therefore main contributors of the pathogenesis of hematopoietic disorders (Burda *et al*., 2010). Despite decades of efforts trying to address how PU.1 and GATA1 negatively regulate each other, little is still known about this mechanism. In this study, we demonstrate that Pu.1 positively regulates *grna* expression, and that this regulatory mechanism is highly conserved in the mammalian granulin. Importantly, our results also indicate that granulin inhibits *gatal* expression *in vivo*. Our findings identify granulin as a previously unrecognized regulator of the Pu.1/Gata1 antagonism and extend previous results on the mechanism of the *gata1* inhibition by Pu.1. Further studies will be required to determine the precise molecular mechanism by which granulin inhibits *gata1* expression. In addition, our first report of the use of CUT&RUN in zebrafish greatly expands our zebrafish genetics toolkit. Since this technique can be performed in a small number of cells with exceptionally low background and high specific signal (Skene & Henikoff, 2017), it will facilitate the discovery of new regulatory networks of TFs that affect gene expression in the zebrafish model, a small animal with relatively low numbers of cells.

In conclusion, with our discovery that granulin is essential for normal myelopoiesis, it is not surprising that its dysregulation leads to pleiotropic effects, since macrophages and neutrophils participate in inflammation, wound healing, tumorigenesis, and neurodegeneration. Our work therefore opens a new field of study that has the potential to impact different aspects of the human health, filling a knowledge gap needed to advance the manipulation of GRN as a therapeutic target. Last, our results are expected to advance understanding of how this protein could be manipulated to treat hematopoietic disorders such as neutropenia or myeloid leukemia. Since granulin is a secreted factor, it may be plausible to be used as a therapeutic target to treat these hematological disorders, expanding the treatment options for these patients.

## Materials and Methods

### Zebrafish husbandry and strains

WT AB* and transgenic zebrafish (*Danio rerio*) embryos and adults were mated, staged, raised and processed as described (Westerfield, 2000) in a circulating aquarium system (Aquaneering) at 28°C and maintained in accordance with ISU and UCSD IACUC guidelines. Mutant *grna^mde54a^* (Solchenberger *et al*., 2015), transgenic *Tg(mpx:eGFP)^i114^* (Renshaw *et al*, 2006), *Tg(gata1:DsRed)^sd2^* (Traver *et al*., 2003), *Tg(mpeg1:eGFP)^gl22^* (Ellett *et al*, 2011), *Tg(kdrl:HsHRAS-mCherry)^s896^* (referred to as *kdrl:mCherry* throughout manuscript) (Bertrand *et al*, 2010), *Tg(Lyz:Dsred2)^nz50^* (Hall *et al*, 2007), *Tg(Myf5:eGFP)^zf42^* (Chen *et al*, 2007), Tg(LCR:eGFP)^cz3325^ (Ganis *et al*, 2012) and various intercrosses of these lines were utilized.

### Morpholino injection

Specific antisense morpholinos (MOs) (Gene Tools) were resuspended in water at 2mM. MOs used in this study were Standard-MO (Gene Tools), pU1-MO (Rhodes *et al*., 2005), irf8-MO (Li *et al*., 2011), Grna-MO1 (also denoted as Grna-MO throughout the manuscript) 5’-TTGAGCAGGTGGATTTGTGAACAGC-3’ (Li *et al*., 2010), Grna control morpholino with 5 nucleotides mismatch 5’-TTGA<u>C</u>CA<u>C</u>GTG<u>C</u>ATTT<u>C</u>T<u>C</u>AACAGC-3’, Grna-MO2 5’-GGAAAGTAAATGATCAGTCCGTGGA-3’ (Li *et al*., 2010), and Grnb-MO 5’-CCACAGCGCAACTCTCACACCTG-3’ (validated in this manuscript). Morpholinos were diluted in water at a concentration of 0.4 mM (Grna-MO), 0.4 mM (Grna mismatch-MO), 0.2 mM (Grna-MO2), 0.6 mM (Grnb-MO), 1.4 mM (irf8-MO) and 2 mM (pu1-MO) with phenol red solution and 2 nl were injected into the yolk ball of one-cell-stage embryos using a micromanipulator (Narishige) and PM 1000 cell microinjector (MDI). For Grnb-MO validation, 20 24 hpf zebrafish embryos injected with Grnb-MO or uninjected controls were collected and RNA was isolated with RNeasy (Qiagen), cDNA generated with qScript Supermix (Quanta BioSciences) and PCR performed with primers Grnb-F2-MO and Grnb-R2-MO (Table S1) for the detection of the wildtype (498 bp) or the mutant (535 bp) *grnb* amplicon. These amplicons were sequenced. The mutant *grnb* amplicon contained 37 extra-nucleotides that changed the reading frame of the *grnb* mRNA.

### Quantitative RT-PCR analysis

RNA was isolated from tissues with RNeasy (Qiagen), and cDNA generated with qScript Supermix (Quanta BioSciences) or iScrip gDNA Clear cDNA Synthesis Kit (BioRad, 1725035). Primers to detect zebrafish transcripts are described in Table S1. qPCR was performed in CFX real-time PCR detection system (BioRad), and relative expression levels of genes were calculated by the following formula: Relative expression= 2^-(Ct[gene of interest]-Ct[housekeeping gene])^.

### Flow cytometry

To quantify the myeloid fraction in kidney marrows from *grna*^-/-^ and *grna*^+/+^ control siblings, adult zebrafish between three to nine months old were anesthetized in tricaine, subjected to cardiocentesis and kidney dissection as previously described (Traver *et al*., 2003). The resulting kidney suspension was gently triturated with a P1000 pipette and filtered with a 30μm cell strainer (Thermo Fisher Scientific, NC9084441) and stained with Sytox Red (Life Technologies) to exclude dead cells. Flow cytometric acquisitions were performed on a LSR-Fortessa (BD) and analyses were performed using FlowJo software (v10.3, Tree Star) as previously described (Traver *et al*., 2003).

To analyze macrophage numbers in Grna morphants, three 48 hpf *Mpeg1:eGFP+* embryos per replicate (three to four replicates per experiment) previously injected with Grna-MO1, Grna-MO2 or Grna mismatch MO were dechorionated with pronase (Roche) and anesthetized in tricaine. Gently triturated with a P1000 pipette and chemically dissociated with liberase TM (Roche) for 20 minutes in agitation. The resulting cell suspension was filtered and stained with Sytox Red to exclude dead cells. Flow cytometric acquisitions were performed on a FACS LSR-Fortessa (Becton Dickinson) and analyses were performed using FlowJo software (v10.3, Tree Star).

### Intracellular flow cytometry

Flow cytometry was performed on a LSR Fortessa (BD) and data were analyzed using FlowJo software (v10.3, Tree Star). Following dissociation of 48 hpf *grna*^+/+^ and *grna*^-/-^ embryos (three embryos per condition) with liberase TM (Roche) as described below, cells were fixed with PFA 4% and permeabilized for 30 minutes on ice with PBS containing 0.1% triton. Intracellular staining was performed with anti zebrafish mfap4 antibody (Gentex, GTX132692), dilution 1:250, followed by staining with 5ug/ml of goat anti-rabbit IgG (H+L) antibody, alexa-488 (Thermo Fisher, A11034).

### Fluorescence-activated cell sorting (FACS)

To isolate embryonic cells for qPCR analysis, approximately 200 16 hpf *Myf5:eGFP+*; 22 hpf *Kdrl1:mCherry+, Gata1:Dsred-;* 48 hpf *Mpx:eGFP;* 48 *Mpeg1:eGFP;* or 36 hpf *LCR:eGFP* embryos were dechorionated with pronase (Roche) and anesthetized in tricaine. Gently triturated with a P1000 pipette. The resulting cell suspension was filtered and stained with Sytox Red to exclude dead cells. Cell sorting of positive fluorescent cells was performed with a FACS ARIAII (Becton Dickinson).

### In Situ Hybridization

WISH was carried out as described (Thisse *et al*, 1993). Probes for the *grna, grnb, pu.1, mfap4, mpx*, and *apoeb* transcripts were generated using the DIG RNA Labeling Kit (Roche Applied Science) from linearized plasmids. Embryos were imaged using a Leica M165C stereomicroscope equipped with a DFC295 color digital camera (Leica) and FireCam software (Leica).

Double fluorescent in situ hybridization (FISH) was performed as previously described (Brend & Holley, 2009) using the following reagents: TSA Plus Cyanine 3/Fluorescein system (NEL753001KT), anti-Digoxigenin-POD, Fab fragments (11207733910), and anti-fluorescein-POD, Fab (11426346910). Embryos were analyzed using a Sp5 confocal (Leica).

### Whole-mount immunohistochemistry

Whole-mount immunohistochemistry for immunofluorescence staining of P-Histone H3 Ser10 in 48 hpf *Tg(Mpx:eGFP)* zebrafish embryos was performed as previously described (Espin *et al*, 2013). The following antibodies were used: rabbit anti-phospho-Histone H3 (Ser10) antibody (Millipore, 06-570) (dilution 1:100), anti-GFP antibody chicken IgY (Aves Labs GFP-1020) (dilution 1:500), goat anti-Chicken IgY (H+L) Alexa-488 (Thermofisher A-11039) (dilution 1:500), donkey anti-rabbit IgG (H+L) Alexa-594 (Thermofisher A21207) (dilution 1:500). The samples were imaged with a Leica Sp5 confocal.

### Enumeration of myeloid progenitors, macrophages and neutrophils

Animals were subjected to WISH for *pu.1* (myeloid progenitors), *mfap4* (macrophages), or *mpx* (neutrophils) at 48 hpf at varying locations (tail or head, listed in figures) and cells expressing the above mentioned transcripts were imaged and manually counted per individual. To image and enumerate macrophages *in vivo* after tail resection, fluorescence microscopy was performed on *Mpeg1:eGFP* transgenic animals. Z-sections of the tail region were captured using Leica Thunder imager with DFC9000 GT camera and LAS X navigator, and manually counted.

### Fin amputation surgeries

48hpf zebrafish embryos were anesthetized with tricaine and a single cut traversing the entire dorsoventral length of the caudal fin was made using a surgical scalpel size #15 (19-200-218, fisher scientific). Embryos were individually isolated and imaged at each given timepoint. Throughout the time course, the initial amputation plane of each embryo was determined by superimposing the image captured at each timepoint with the image of the initial cut using Adobe Photoshop CS6. The caudal-most tip of notochord was used as a landmark for spatial alignment. Quantification of regenerate area was determined using ImageJ.

### Cytology

Cytospin preparations were made with 1×10^5^ to 2×10^5^ kidney cells cytocentrifuged at 300 rpm for 5 min onto glass slides in a Thermo Scientific Shandon CytoSpin 4. Cytospin preparations were processed through Wright-Giemsa stains (Fisher Scientific, 5029782) for morphological analyses and differential cell counts. Briefly: Wright Stain Solution was placed upon the smear for 2 mins and washed with distilled water. Subsequently, the slides were placed for 10 minutes in a coplin jar containing Giemsa and rinsed with distilled water.

### Morphological analyses and differential cell counts of kidney marrow hematopoietic cells

200 non-erythroid nucleated differential cell counts were assessed in *grna*^-/-^ or control *grna+/+* siblings kidneys after cardiocentesis, kidney dissection, cytospin and staining with Wright-Giemsa as described here. The morphological features to identify each cell type are the following. Early myeloid precursors, round to ovoid in shape with a round to ovoid-shaped pink nucleus. Chromatin pattern finely granular and semi frequently with distinct round nucleoli. High nuclei to cytoplasmic (N:C) ratio with a moderate amount of deep blue cytoplasm. Late myeloid precursors (immature neutrophils): round to ovoid to bilobed nuclei are with light purple to darker pink coloration. Slightly clumped chromatin pattern and no distinct nucleoli. Moderate N:C ratio with a blue to light blue cytoplasm. Neutrophils (mature): smaller in size than late myeloid precursors. Round to ovoid to bilobed to segmented dark purple nucleus smaller than late myeloid precursors. Moderate N:C ratio with a light blue cytoplasm and often bubbly vacuolated appearance. Macrophages: round to ovoid in shape. Nuclear shapes included round, ovoid, bilobed, and lobulated forms that have a pale pink to purple coloration. The chromatin pattern is finely stippled that occasionally has distinct nucleoli. Moderate N:C ratio that is moderately high with a blue appearance frequently vacuolated and sometimes contained melanin pigment. Lymphocytes: small round to ovoid-shaped cells. Round to ovoid-shaped nucleus, coarsely clumped chromatin pattern, and no distinct nucleoli. High N:C ratio with a scant amount of pale blue cytoplasm.

### CUT&RUN

Adult wild type AB* zebrafish between three to nine months old were anesthetized in tricaine, subjected to cardiocentesis and kidney dissection as previously described (Traver *et al*., 2003). The resulting kidney suspension was gently triturated with a P1000 pipette and filtered with a 30μm cell strainer (Thermo Fisher Scientific, NC9084441). 90,000 −120,000 cells were used per condition to perform CUT&RUN using CUT&RUN assay kit (86652S, Cell signaling) with an anti zebrafish Pu.1 antibody (GTX128266, GeneTex) (4 μl) or rabbit (DA1E) IgG Isotype control antibody (as recommended in kit protocol) following the manufacturer instructions with the following modifications: Digitonin dilution 5:1000. Incubation time for primary antibody: 10 hours at +4°C; for pAGMase: 1 hour at +4°C. Digestion time: 30 min.

### *In silico* prediction of Pu.1 binding sites

To identify potential binding sites for Pu.1 in the promoter and enhancer regions of *grna*, the entire *grna* gene sequence plus 10Kb 5’ upstream *grna* were subjected to Pu.1 binding sites prediction using Consite (http://consite.genereg.net/) and Tomtom (http://meme-suite.org/tools/tomtom). The transcription factor binding profile matrix for Pu.1 ID: MA0080.4 from JASPAR was utilized when using Tomtom, and user defined profile (http://jaspar.genereg.net/matrix/MA0080.4/) and 70% identity analyses were used for Consite. The fragment located at +23765 from the *grna* ATG start codon was used as a control, since it lacked any predicted binding sites for Pu.1.

### Preparation of RNA for RNA sequencing

Adult *grna*^-/-^ and *grna*^+/+^ control fish (three fish per condition) were subjected to cardiocentesis and kidney dissection as described here. RNA was isolated with RNeasy (Qiagen) following the manufacturer instructions. Total RNA was assessed for quality using an Agilent Tapestation 4200, and samples with an RNA Integrity Number (RIN) greater than 8.0 were used to generate RNA sequencing libraries using the TruSeq Stranded mRNA Sample Prep (Illumina, San Diego, CA). Samples were processed following manufacturer’s instructions, starting with 50 ng of RNA and modifying RNA shear time to five minutes. Resulting libraries were multiplexed and sequenced with 75 basepair (bp) single reads (SR75) to a depth of approximately 20 million reads per sample on an Illumina HiSeq 4000. Samples were demuxltiplexed using bcl2fastq v2.20 Conversion Software (Illumina, San Diego, CA).

### Processing of RNA-seq dataset

RNA-seq data was mapped to Reference Consortium Zebrafish Build 10 (UCSC Genome GRC10/danRer10; Sept 2014) using Olego (Wu *et al*, 2013) and normalized using standard analysis pipelined such as cufflinks (Trapnell *et al*, 2009; Trapnell *et al*, 2012; Trapnell *et al*, 2010). feautreCounts (Liao *et al*, 2014) from subread package is used to compute the raw read counts for each gene. TPM (Transcripts Per Millions) (Li & Dewey, 2011; Pachter, 2011) values were computed from the raw read counts using a custom perl script and log2(TPM+1) is used to compute the final log-reduced expression values. DESeq2 1.26.0 (Love *et al*, 2014) R package is used to compute differentially expressed genes at 1% false discovery rate. Volcano plot and heatmap were created using python matplotlib package (version 2.1.1).

### Statistical analysis

Data were analyzed by unpaired T-test in GraphPad Prism 8. In all figures, solid red bars denote the mean, and error bars represent S.E.M. * p< 0.05, ** p < 0.01, *** p < 0.001, **** p < 0.0001, n.s. not significant, n.d. not detected.

## Acknowledgments

This work was supported by the NIH-NIDDK K01 (7K01DK115661) and R03 (R03DK125661) awards, Iowa State University start-up funds, postdoctoral fellowship from Fundacion Seneca, Agencia de Ciencia y Tecnologia de la Region de Murcia, and American Heart Association (16POST30690005) to R.E-P. We thank Roy J. Carver Charitable Trust for the zebrafish research facility in the Advanced Teaching and Research Building at ISU. We thank Karen Ong for technical assistance, and Jesus Olvera and Cody Fine of the UCSD Human Embryonic Stem Cell Core Facility for technical assistance of flow cytometry experiments. This work was made possible in part by the CIRM Major Facilities grant (FA1-00607) to the Sanford Consortium for Regenerative Medicine. We are grateful to Kristen Jepsen, Eugenia Ricciardelli, and Stephanie Hadimulia of the Institute of Genomics Medicine (IGM) at UCSD for technical assistance of RNA-sequencing experiments. This publication includes data generated at the UC San Diego IGM Genomics Center utilizing an Illumina NovaSeq 6000 that was purchased with funding from a National Institutes of Health SIG grant (#S10 OD026929).

## Authorship contributions

R.E.-P, C.C, O.F, D.T. designed experiments; R.E.-P, C.C., O.F., E.S, X.C, L.L, A.M, B.S, M.M., performed research; R.E.-P, C.C, X.C, O.F., E.S, L.L, A.M, B.S, D.S, M.M., D.T. analyzed data; and R.E.-P., C.C., and D.T. wrote the paper with minor contributions from remaining authors.

---

**Supplementary Figure 1:**
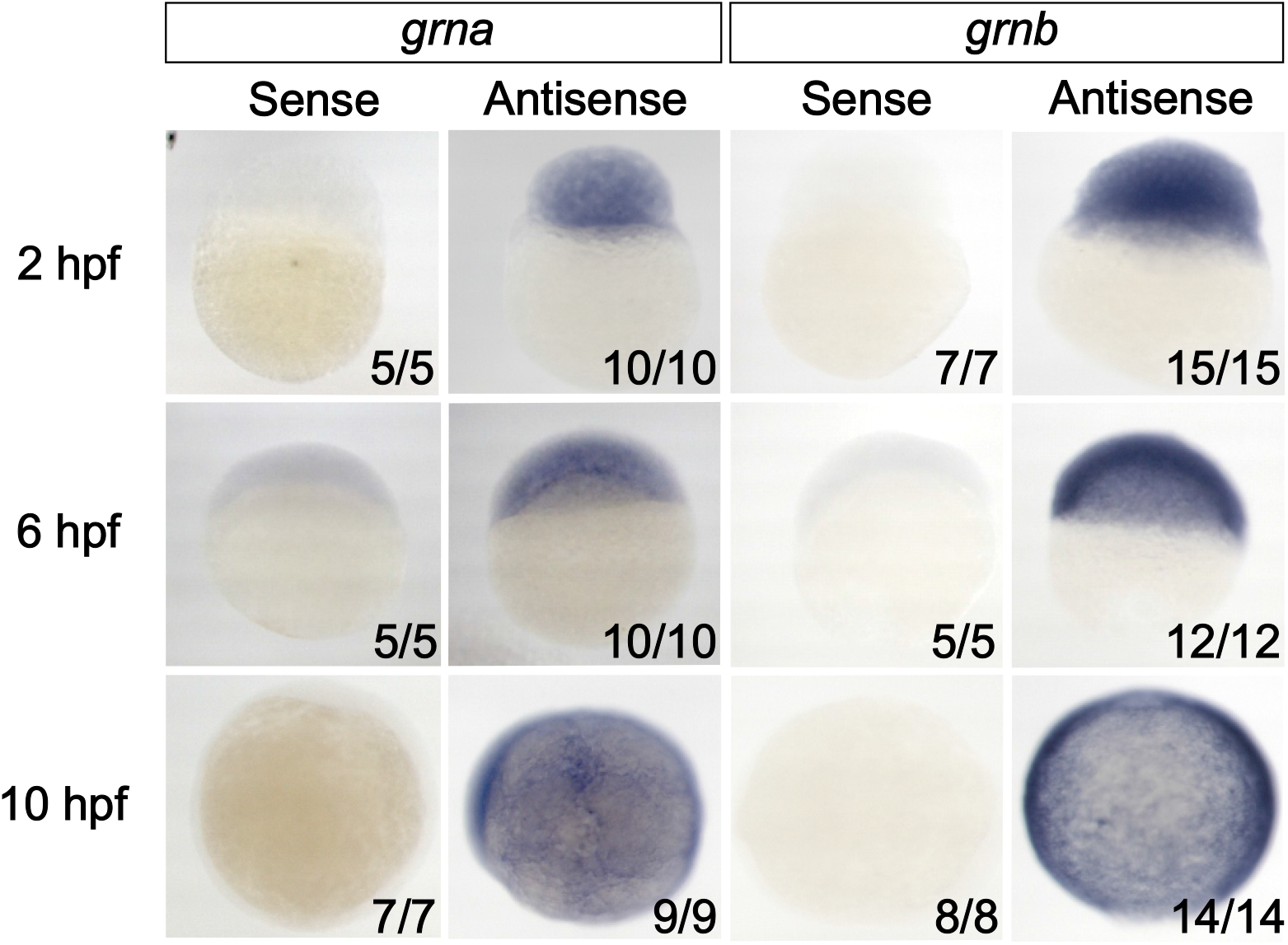
*grna* and *grnb* transcripts are ubiquitously expressed during early zebrafish embryonic development. WISH of 2, 6 and 10 hpf AB* zebrafish embryos hybridized with *grna* antisense and control sense probes (left panel), or *grnb* antisense and control sense probes (right panel). Numbers represent embryos with indicated expression.

**Supplementary Figure 2:**
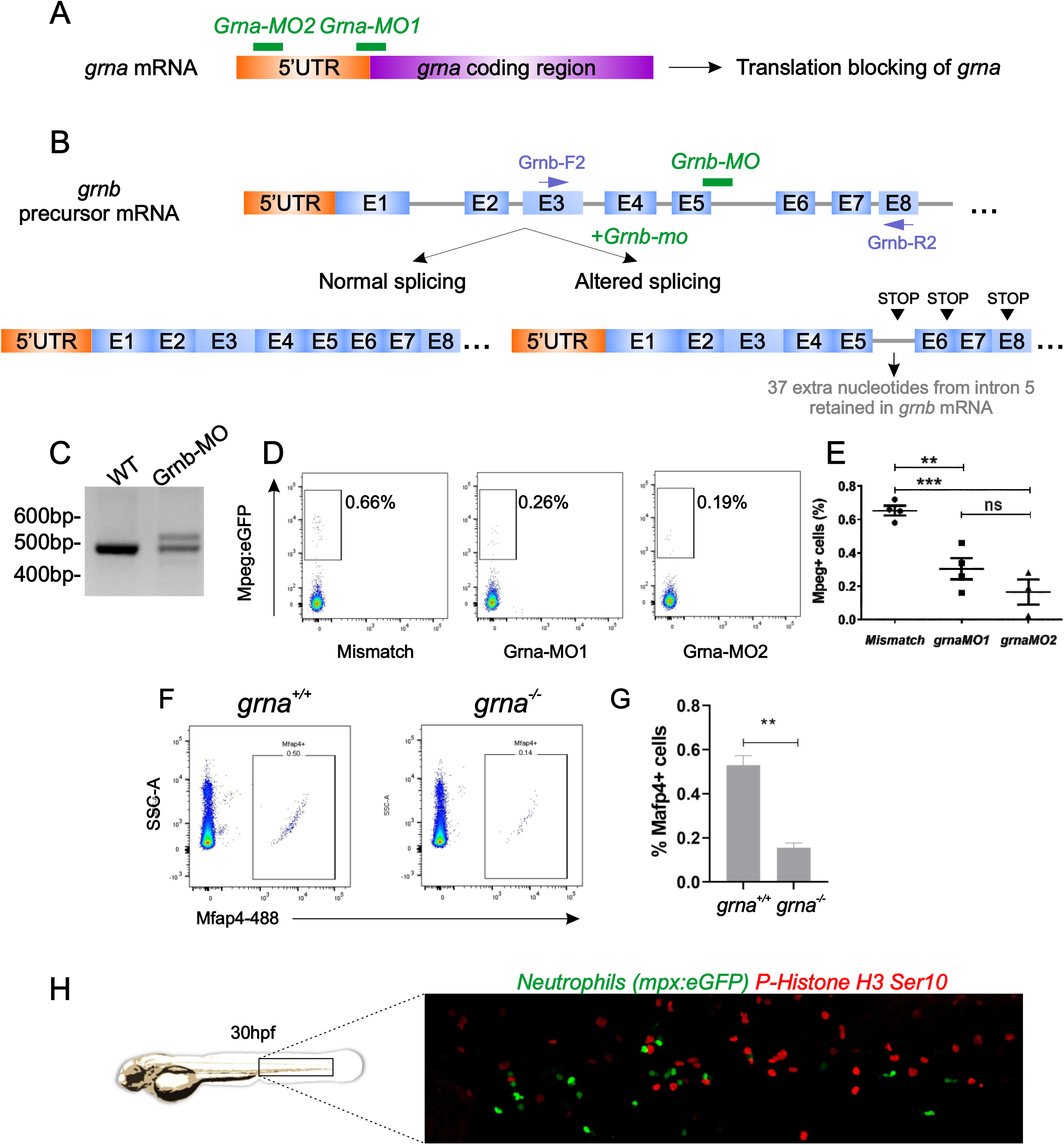
*grna* knockdown impairs macrophage development. **(A-B)** Schematic illustration of the *grna* and *grnb* knock-out and strategies. **(A)** Grna-MO1 and Grna-MO2 are 5’UTR-ATG translation blocking morpholinos. **(B-C)** Efficiency of splice-blocking MO against zebrafish *grnb*. RT-PCR analysis of WT and Grnb-MO injected embryos induced altered splicing of the *grnb* transcripts at 24 hpf. A 535 bp product containing 37 nucleotides insertion from *grnb* intron 5 (upper band) was detected in contrast to a 498 bp control (lower band in Grnb-MO injected embryos and WT control). This leads to an aberrant *grnb* mature mRNA with a shift in reading frame, which results in premature stop codons along the *grnb* mRNA. The annealing of MOs (green lines) and the inframe premature stop codons (arrowheads) are indicated. **(D)** Representative dots plots of 36hpf *Mpeg1.1:eGFP* embryos injected with mismatch, Grna-MO1 and Grna-MO2. **(E)** Quantification of Mpeg+ macrophages in Mismatch control n=4, Grna-MO1 n=4, and Grna-MO2, n=3 embryos. Each dot represents the percentage of Mpeg+ cells from the total events from 3 embryos. Horizontal lines and error bars indicate means ± SEM (**p<0.01, ***p<0.001, ns: no significant). **(F-G)** Representative dot plot of intracellular flow cytometry using a specific antibody against the macrophage marker Mfap4 at 48 hpf grna^+/+^ (left) or grna^-/-^ embryos (right). Quantification shown in **(G)**. **(H)** WIHC for eGFP (green) and phospho-Histone 3 (Ser10) (red) in the caudal hematopoietic tissue of 30 hpf *Mpx:eGFP* embryos shows lack of colocalization between both markers.

**Supplementary Figure 3:**
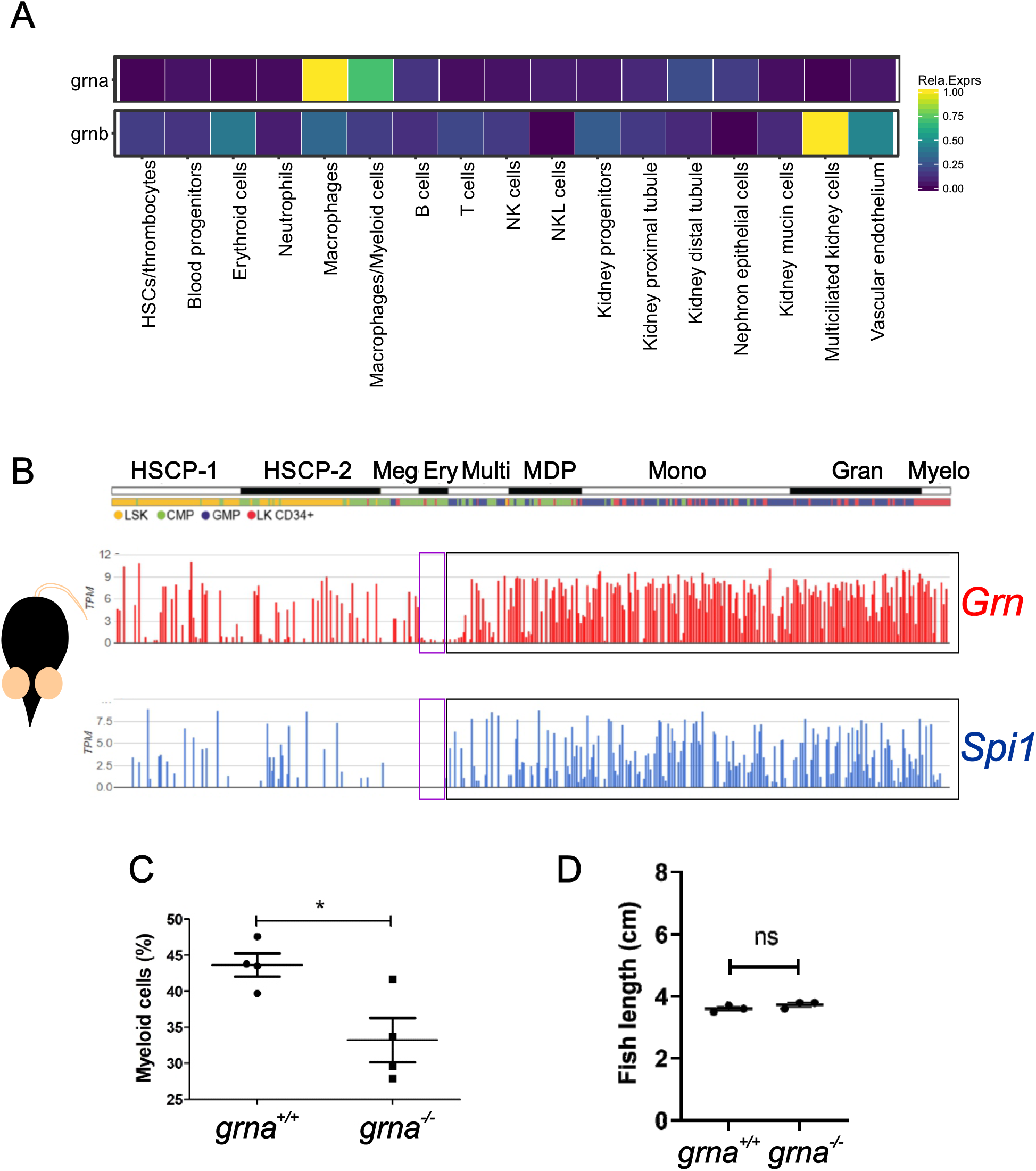
Conserved gene expression levels among zebrafish *grna* and mouse *Grn* in hematopoietic cells. **(A)** Heat map derived from the online visualizer “Single Cell inDrops RNA-Seq Visualization of Adult Zebrafish Whole Kidney Marrow” (https://molpath.shinyapps.io/zebrafishblood/#pltly) (Tang *et al*., 2017) for *grna* and *grnb* showing low or inexistent expression of *grnb* by kidney marrow hematopoietic cells and high expression by multiciliated kidney cells and vascular endothelium. In contrast, *grna* is highly expressed by macrophages and myeloid cells. **(B)** Gene expression levels in Transcripts Per Kilobase Million (TPM) for Granulin (*Grn*, red) and Pu.1 (*Spi1*, blue) from single cell RNA-seq data of mouse hematopoietic cells (Olsson *et al*., 2016) showing high correlation between *Grn* and *Spi1* expression. **(C)** Quantification of the percentage of myeloid cells gated in Figure 3C in *grna*^+/+^ (n=4) and *grna*^-/-^ (n=4) zebrafish kidney marrows. **(D)** Quantification of zebrafish body length of *grna*^-/-^ and *grna*^+/+^ control siblings. Horizontal lines and error bars indicate means ± SEM. Ns, not significant; *p < 0.05. HSCP, Haematopoietic Stem Cell Progenitor; Meg, megakaryocytic; Ery, erythrocytic; Multi, Multi-lineage primed; MDP, monocyte-dendritic cell precursor; Mono, monocytic; Gran, granulocytic; Myelo, myelocyte (myelocytes and metamyelocytes).

**Supplementary Figure 4:**
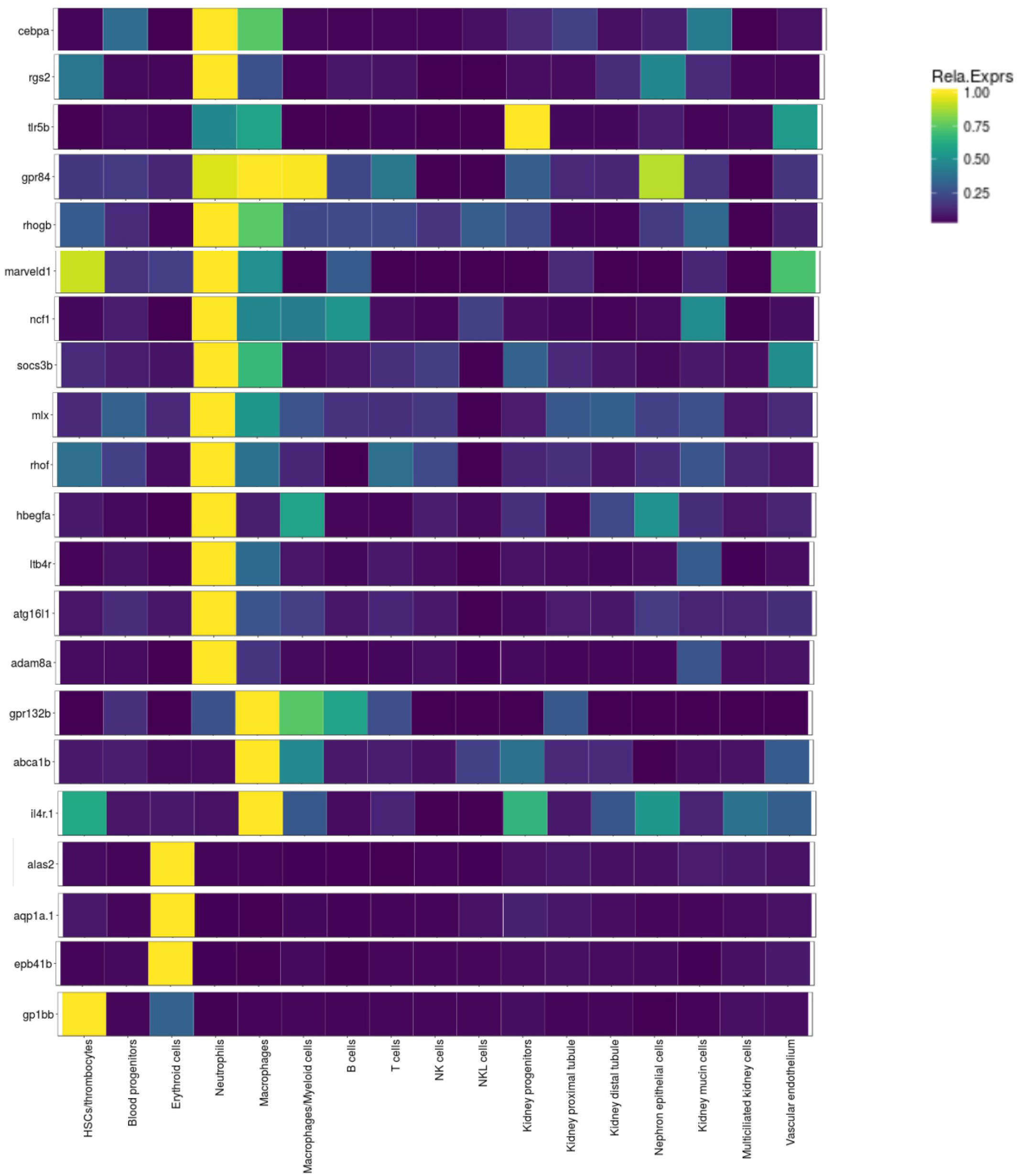
Grna depletion leads to decreased expression of myeloid specific genes and increased erythroid genes in the kidney marrow. Heat map of the relative gene expression derived from the online visualizer “Single Cell inDrops RNA-Seq Visualization of Adult Zebrafish Whole Kidney Marrow” (https://molpath.shinyapps.io/zebrafishblood/#pltly) (Tang *et al*., 2017) of genes significantly enriched **(A)** or depleted **(B)** from RNA-seq of *grna*^-/-^ versus *grna*^+/+^ kidney marrows.

**Supplementary Figure 5:**
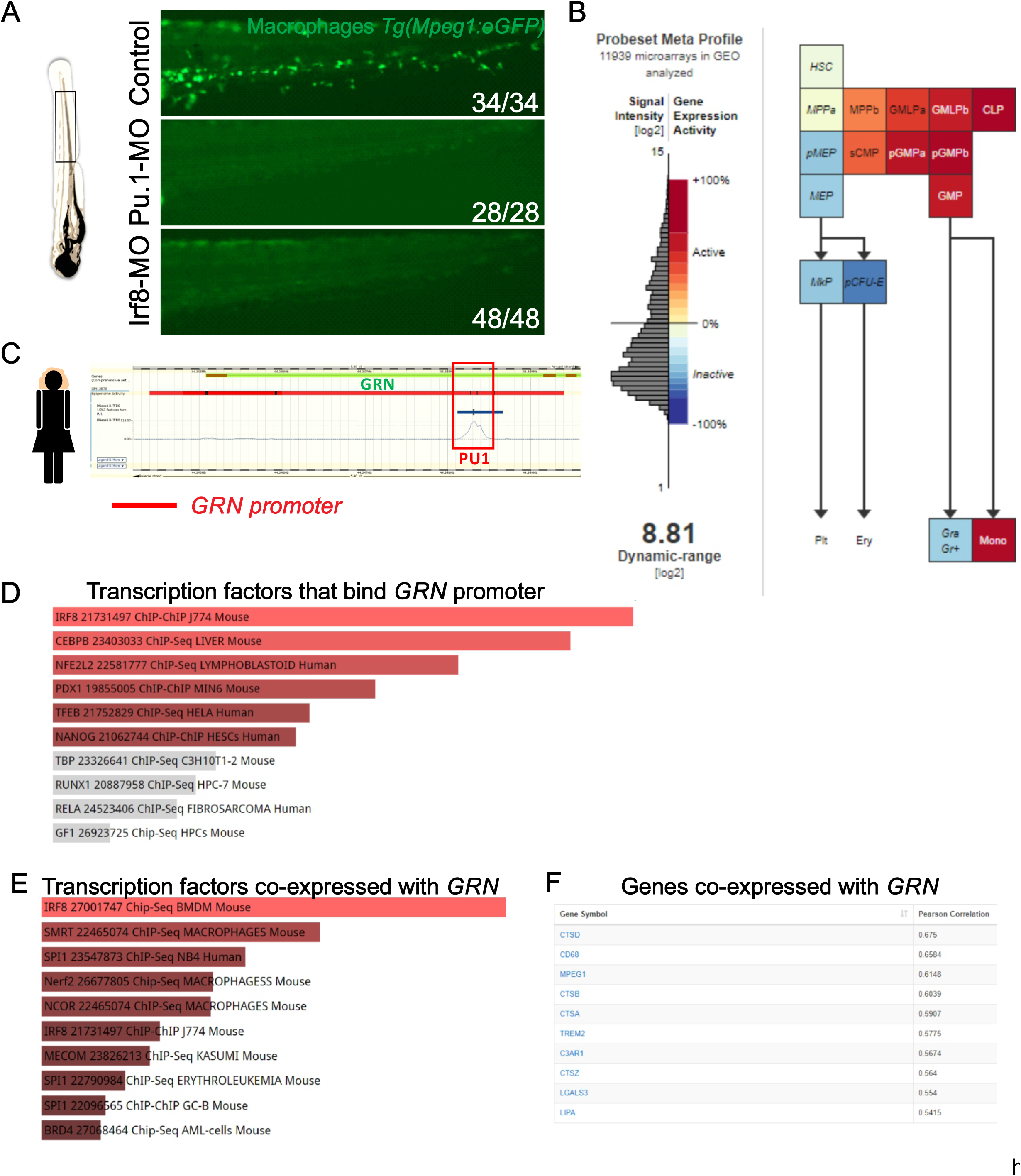
Conserved regulatory expression of granulin in mammals and zebrafish. **(A)** *Mpeg1:eGFP* transgenic embryos were injected with control (Std), Pu.1, or Irf8 MOs and the tail region was visualized by fluorescence microscopy at 48 hpf. **(B)** Mouse hematopoietic model showing the dynamic expression of *Irf8* derived from microarray data (Affymetrix Mouse Genome 430 2.0 Array). Notice that lymphocyte differentiation beyond CLP is not shown here. **(C)** Screenshot of the regulatory feature of *ensembl.org* showing that PU.1 binds the *GRN* promoter in human hematopoietic cell lines based on epigenetic marks from experimental data available through *ensembl.org*. **(D)** Screenshot of mammalian granulin queried by ChEA showing the transcription factors whose peaks were detected at the granulin promoter. TFs were ranked based on p-value (red colors indicate p<0,05; gray colors indicate p>0,05). **(E)** Results from Harmonizome combined with ChEA showing the TFs that were co-expressed with the mammalian granulin. TFs were ranked based on p-value (red colors indicate p<0,05). **(F)** Screenshot of mammalian granulin queried by Harmonizome (https://amp.pharm.mssm.edu/archs4/gene/GRN#correlation) showing the first 10 most similar genes based on co-expression ranked by descendent Pearson correlation. HSC, Hematopoietic Stem Cell; MPP, Hematopoietic multipotential progenitors; GMLP, granulocyte–monocyte–lymphoid progenitor; CLP, common lymphoid progenitor; pMEP, pre of megakaryocyte-erythroid progenitor; MEP, megakaryocyte-erythroid progenitor; MkP, Megakaryocyte progenitor; PCFU-e, Colony Forming Unit-Erythroid; Plt, platelets; Ery, Erythrocytes; sCMP, strict common myeloid progenitor; pGMP, pre-granulocyte/macrophage; GMP, granulocyte/macrophage progenitors; Gra Gr+, granulocytes; Mono, monocytes.

**Table S1:**
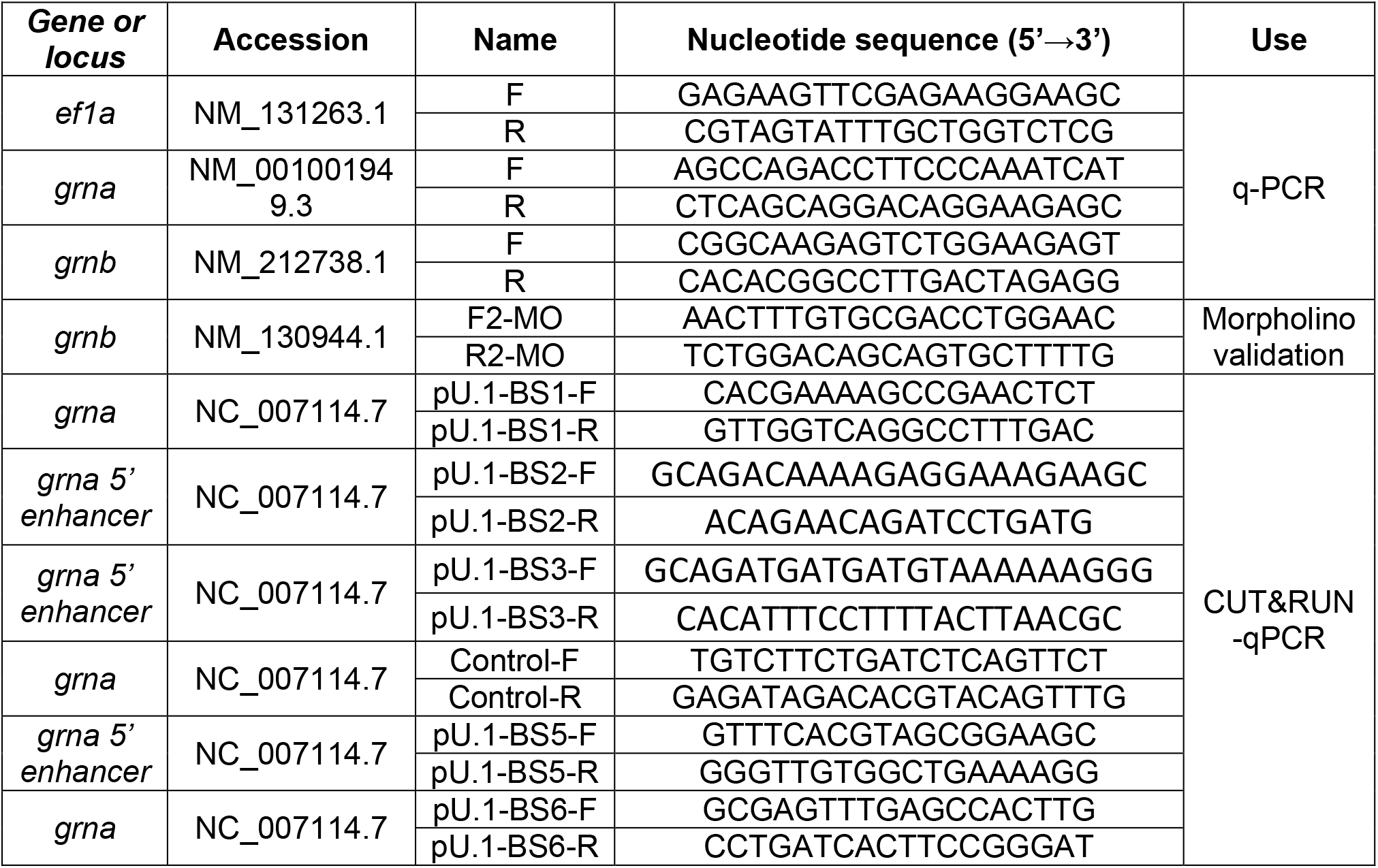
Primers used in this manuscript.

